# Nitrogen competition is the general mechanism underlying cnidarian-Symbiodiniaceae symbioses

**DOI:** 10.1101/2022.06.30.498212

**Authors:** Guoxin Cui, Jianing Mi, Alessandro Moret, Huawen Zhong, Shiou-Han Hung, Salim Al-Babili, Manuel Aranda

## Abstract

Symbiotic associations with Symbiodiniaceae have evolved independently across a diverse range of cnidarian taxa including reef-building corals, anemones and jellyfish, yet the molecular mechanisms underlying their regulation and repeated evolution are still elusive. Here we show that despite their independent evolution, cnidarian hosts employ the same mechanism of symbiont control in which symbiont-derived glucose is used to assimilate nitrogenous waste via amino acid biosynthesis to limit the availability of nitrogen to the symbionts. In this metabolic interaction, glucose significantly reduces symbiont density while ammonium promotes symbiont proliferation. We show that glucose-derived ^13^C and ammonium-derived ^15^N are co-incorporated into amino acids by the hosts. Metabolic differences between the hosts further suggest that corals are more susceptible to environmental stress and symbiosis breakdown due to their increased energy demands to satisfy calcification. Our results reveal the general metabolic interaction underlying these symbioses and provide a parsimonious explanation for their repeated evolution.

## Introduction

The mutualistic symbiotic relationship between marine invertebrates and dinoflagellates in the family Symbiodiniaceae is one of the most common eukaryote-eukaryote endosymbiosis in our oceans and fundamental to coral reef ecosystems (Stat et al., 2006). The symbiotic association with Symbiodiniaceae provides the hosts with photosynthetically derived carbohydrates and allows them to thrive in the oligotrophic environments of tropical oceans. Symbiodiniaceae symbioses have evolved convergently across a broad range of marine phyla, including single-celled foraminifera, sponges, cnidarians, platyhelminths, and mollusks (LaJeunesse et al., 2018; Melo Clavijo et al., 2018). Among these phyla, cnidarians have arguably evolved the largest diversity in Symbiodiniaceae symbioses. Two out of the four cnidarian subphyla that diverged >700 Mya, Anthozoa and Scyphozoa, have species that evolved symbiotic relationships with Symbiodiniaceae (Colley and Trench, 1983; Davy et al., 2012; Furla et al., 2005). However, anthozoans, which include octocorals, anemones, and reef-building corals, among others, have evolved the highest diversity in Symbiodiniaceae relationships. Of all the symbiotic cnidarians, reef-building corals are the best-studied due to their economical and ecological importance (Stat et al., 2008). The evolution of symbiosis turned the ancestor of corals into the ecosystem founders they are today by enabling them to build the framework of one of the most productive and biodiverse ecosystems on our planet, coral reefs. Unraveling the molecular mechanisms underpinning these relationships will not only provide valuable insight into the mechanisms underlying the regulation of these associations and their convergent evolution but also provide critical information for our understanding of its stress-related breakdown known as bleaching.

## Results

### Mechanisms of host-symbiont metabolic interactions

The repeated evolution of these symbioses across such a diverse range of phyla suggests that a common mechanism might exist, which regulates the interactions between hosts and symbionts. These interactions need to allow for bidirectional nutrient exchange to establish an environment conducive to symbiont growth and function, but at the same time provide a mechanism to regulate and control symbiont proliferation within host tissues. Several possible mechanisms have been proposed and investigated in the past, including specific protein machinery that allows the host to directly interfere with the symbiont cell cycle (Dimond et al., 2013; Tivey et al., 2020), to host-controlled preferential expulsion of dividing symbionts (Baghdasarian and Muscatine, 2000), as well as more simple nutrient-flux-based models (Cui et al., 2019; Smith and Muscatine, 1999). However, the repeated evolution of these relationships across such a vast range of taxonomic groups suggests that the establishment of these symbioses might not require the *de novo* evolution of complex protein machinery. Hence, simple models requiring less evolutionary novelties should be considered more likely. Previous studies have suggested that symbiont proliferation might indeed be controlled via the limitation of essential nutrients such as nitrogen (Cui et al., 2022; Cui et al., 2019; Falkowski et al., 1993; Wang and Douglas, 1998; Xiang et al., 2020). Based on these findings, we proposed a simple metabolic model in which the host uses the photosynthesis-derived sugar provided by the symbionts to assimilate its own waste nitrogen via the GS/GOGAT cycle and subsequently incorporates it into non-essential amino acids [**Figure 1**, see also Cui et al. (2019)]. The model is based on a simple metabolic interaction that allows the host to convert sugar and waste nitrogen into valuable amino acids while simultaneously providing a mechanism for symbiont control without the requirement of additional means to regulate symbiont cell numbers.

**Figure 1.**
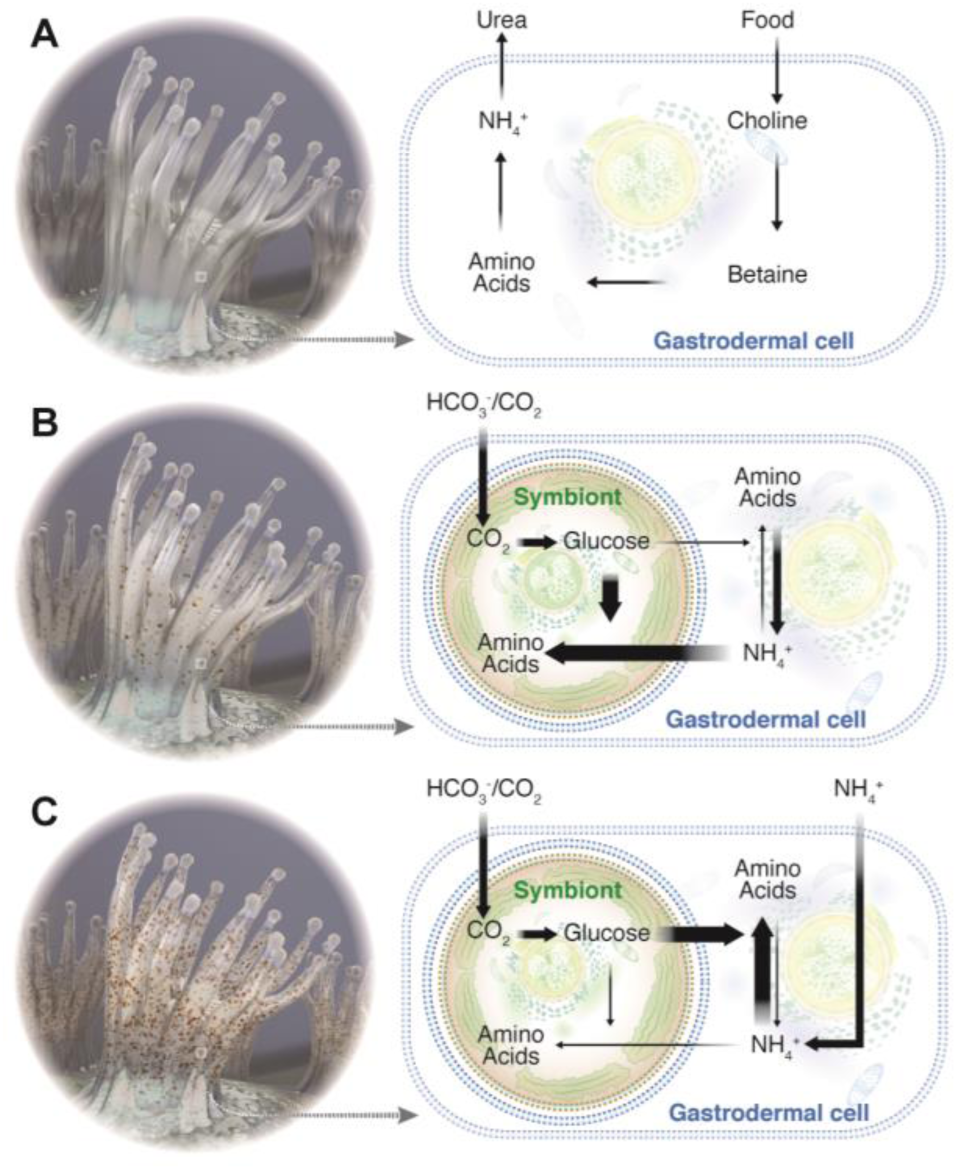
Nutrient-flux-based negative feedback mechanism underlying symbiont population control. (**A**) At the aposymbiotic state, animal hosts are limited by the availability of energy-rich carbohydrates. They take in organic carbon from food and release nitrogenous waste to the surrounding environment. (**B**) At the early stages of symbiosis, the host experiences an increasing provision of energy-rich photosynthates from the symbionts and gradually starts shifting towards a nitrogen-limited state. (**C**) At the fully symbiotic state, symbiont-provided glucose increases the ability of the host to assimilate its own waste nitrogen, which leads to a further reduction in nitrogen availability to the symbionts and results in a further decrease of symbiont proliferation rates that eventually reaches a balance where the symbiont population is stable.

### Glucose and ammonium modulate symbiont density

To determine if this model represents the general mechanism underlying the symbioses across different cnidarians, we tested it in three distantly related species that evolved symbiosis independently, the reef-building coral *Stylophora pistillata*, the sea anemone *Exaiptasia diaphana*, and the upside-down jellyfish *Cassiopea andromeda*. These species are representatives of three different cnidarian classes that evolutionary diverged >700 Mya ago (Park et al., 2012; Wang et al., 2021).

Based on our proposed metabolic model (**Figure 1**), we hypothesized that symbiont cell density in symbiotic hosts is tightly controlled through a negative feedback response driven by the availability of glucose and ammonium. In this self-regulating system, increasing glucose levels are expected to promote the capacity of the host to assimilate ammonium and, thus, to limit the availability of nitrogen to the symbionts. This reduction in nitrogen availability is expected to result in a decrease of the symbiont cell density over time since the reduced nitrogen level would not be sufficient to support the original number of symbionts. Conversely, the symbiont cell density is expected to increase when the availability of ammonium in the system increases.

To test this hypothesis in the three selected cnidarian species, we manipulated the levels of glucose and ammonium in their surrounding environment and analyzed their responses on the level of symbiont density. As predicted by our model, the supplementation with glucose resulted in significantly lower symbiont cell densities in the reef-building coral *Stylophora pistillata* (**Figure 2A**, unpaired two-tailed *t* test, *p* = 0.000003), the sea anemone *Exaiptasia diaphana* (**Figure 2B**, *p* = 0.008), and the upside-down jellyfish *Cassiopea andromeda* (**Figure 2C**, *p* = 0.039). Conversely, symbiont cell densities increased significantly in *S. pistillata* (*p* = 0.002) and *E. diaphana* (*p* = 0.025), when ammonium was supplied, while *C. andromeda* showed an increasing, but nonsignificant trend (*p* = 0.382) (**Figure 2**). Interestingly, experiments using both glucose and ammonium combined did not show the reduction in symbiont density observed in response to glucose alone (**Figures S1 and S2**). This finding confirms that the observed reduction in symbiont density in response to glucose provision is indeed a direct response and not an unspecific stress response that induces bleaching. To provide additional confirmation of these responses in corals, we repeated these experiments in the coral *Acropora hemprichii* which showed the same responses as *S. pistillata* (**Figure S2**).

**Figure 2.**
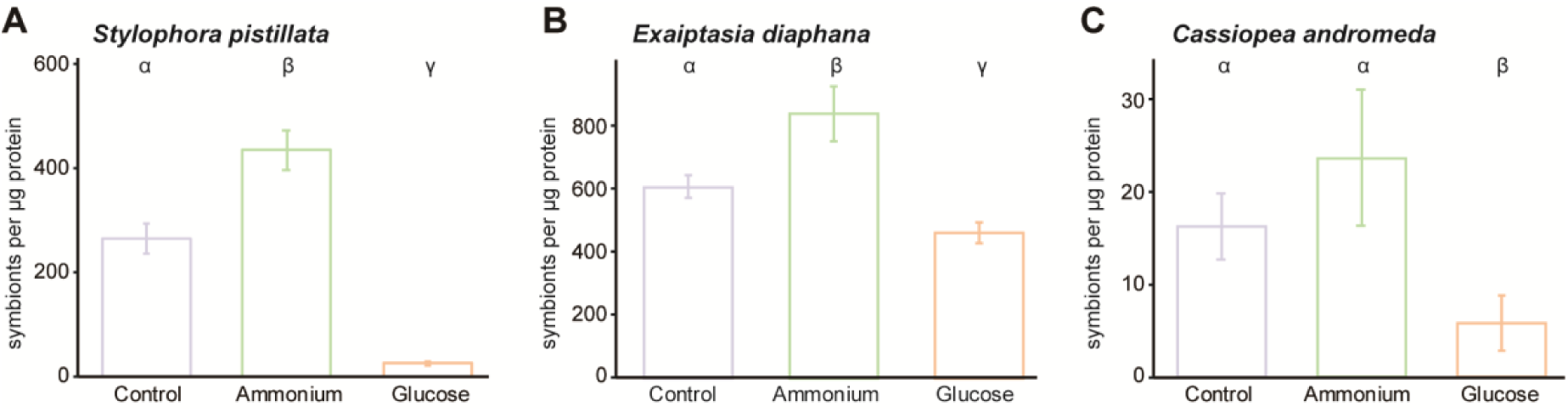
Symbiont cell density changes induced by the availability of glucose and ammonium. Symbiont cell densities were calculated by normalizing total symbiont numbers to host protein content. Bars represent the standard error of the mean. Greek letters indicate statistical differences with a significance cut-off at *p* = 0.05.

### Host-dependent ammonium assimilation and amino acid synthesis

The observed changes in symbiont cell densities in response to the availability of glucose and ammonium aligned with the predictions based on our metabolic model. Hence, we further hypothesized that host-driven ammonium assimilation via amino acid biosynthesis is the molecular pivot underlying symbiont population control. To evaluate this hypothesis, we performed stable isotope tracer analysis using ^13^C labeled glucose and ^15^N labeled ammonium by ultra-high-performance liquid chromatography-high resolution mass spectrometry (UHPLC-HR-MS).

Using UHPLC-HR-MS, we examined the isotopic profiles of metabolic intermediates of the GS/GOGAT and amino acid biosynthesis pathways from host animals supplemented with ^13^C_6_-glucose, ^15^N-ammonium, and combined ^13^C_6_-glucose and ^15^N-ammonium. The high-resolution mass spectra acquired with enhanced resolution to 280,000 m/Δm (at *m/z* = 200), facilitated unambiguous identification and distinction of targeted compounds with different stable ^13^C and ^15^N isotopic compositions. Here we present the identification of glutamine as an example (**Figure 3A**). The natural isotopic distribution of glutamine standard shows two clear isotopic ions increasing by 1 amu, which are recognized as [^13^CC_4_H_10_N_2_O_3_]^+^ ion at *m/z* 148.07948 and [C_5_H_10_^15^NNO_3_]^+^ ion at *m/z* 148.07321, respectively. In addition, their intensities are about 5% and 1% of the monoisotopic ion [C_5_H_10_N_2_O_3_]^+^, respectively. Compared to glutamine standard, the mass spectrum of endogenous glutamine of *E. diaphana* incubated with both ^13^C_6_-glucose and ^15^N-ammonium indicates the presence of various glutamine molecule compositions containing different amounts of ^13^C and ^15^N atoms (**Figure 3A**). Following this strategy, we profiled metabolites including 3-phosphohydroxypyruvate, glutamate, glutamine, O-phospho-*L*-serine, serine, and glycine, which are associated with GS/GOGAT-mediated ammonium assimilation and subsequent amino acid biosynthesis in *S. pistillata*, *E. diaphana*, and *C. andromeda* (**Figures 3B, S3-S25**).

**Figure 3.**
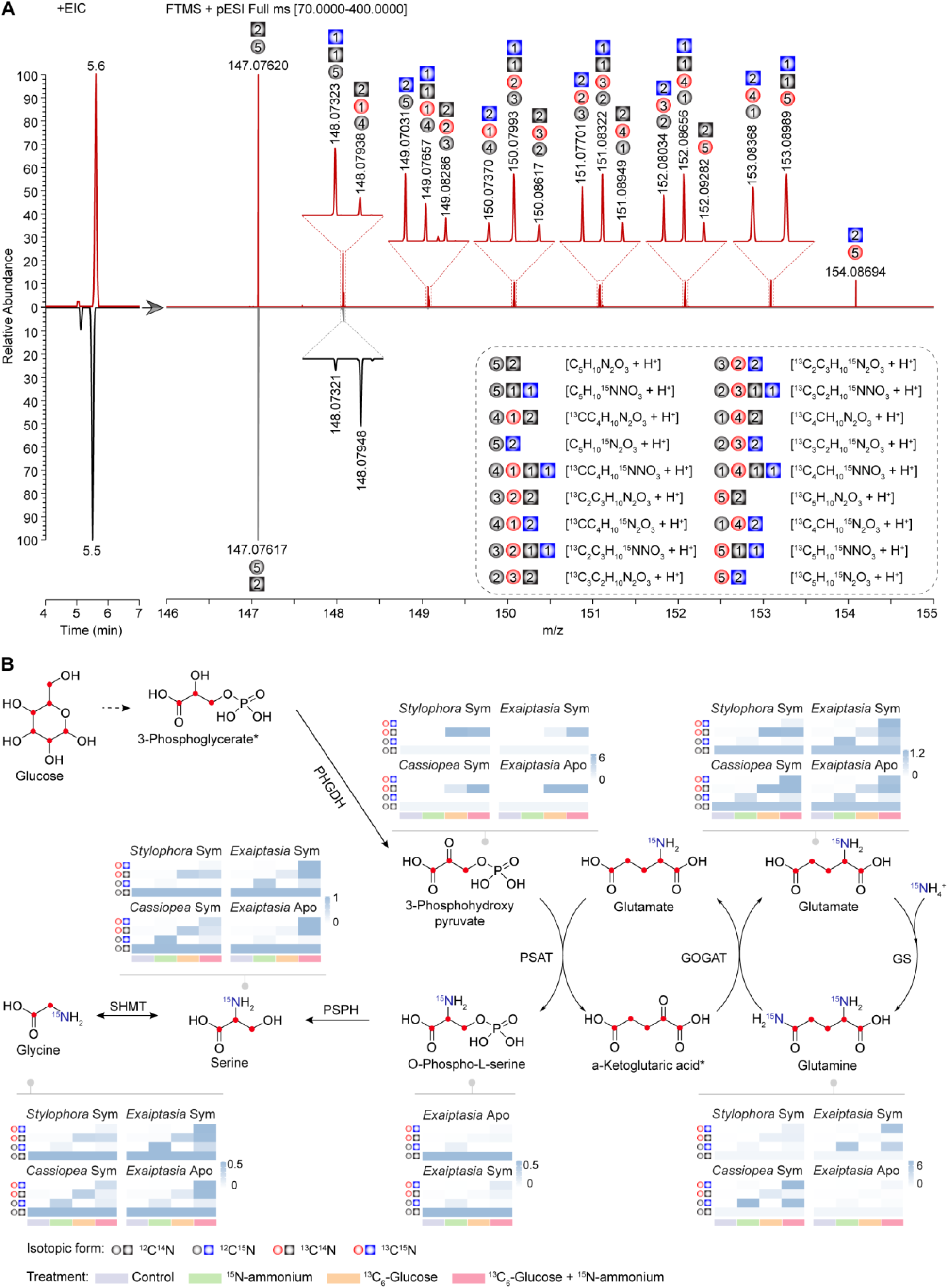
Identification of isotope-labeled metabolites from *Aiptasia* fed with ^13^C_6_-glucose and ^15^N-ammonium using UHPLC-HR-MS. (**A**) Extracted ion chromatograms (EIC, Left) and the isotopic distributions (Right) of glutamine from *E. diaphana* incubated with ^13^C_6_-glucose and ^15^N-ammonium (Upper) and the corresponding glutamine standard (Down). The inset corresponds to a zoom of the area in which different isotopologue compositions of glutamine (dashed box) were identified using HR-MS. The gray ball and square indicate ^12^C atom and ^14^N atom, respectively; the red ball indicates ^13^C atoms, the blue square indicates ^15^N, and the number of carbon and nitrogen atoms are inserted in the corresponding shapes. (**B**) Metabolic footprinting of stable isotopes in the three selected cnidarian species. The proposed ^13^C and ^15^N isotope labelings are indicated as red dots or written in blue color in the structural formulas. Heatmap color indicates the relative abundance of isotope-labeled metabolites normalized to their natural non-labeled forms. Sym, symbiotic state; Apo, aposymbiotic state; *, undetectable metabolite.

For further comparison of each metabolite with different nitrogen and carbon isotopic compositions from the different supplementation experiments, we first normalized the levels of isotopic ions to the abundance of the natural monoisotopic ion and then summarized them according to their isotopic compositions (^12^C^14^N, ^13^C^14^N, ^12^C^15^N, and ^13^C^15^N).

In animals supplemented with ^13^C_6_-glucose, the proportion of ^13^C-containing 3-phosphohydroxypyruvate, one of the intermediate metabolites derived from glycolysis, increased dramatically from the natural level (< 5%) to >50% (mean percentage ± SE, 86.89% ± 0.88% in *S. pistillata*, 53.43% ± 6.97% in *E. diaphana*, and 69.75% ± 2.46% in *C. andromeda*). Similar isotopic results were observed in downstream metabolic intermediates of the amino acid biosynthesis pathway via the GS/GOGAT cycle (**Figure 3B**).

The provision of ^15^N-ammonium in combination with ^13^C_6_-glucose further increased the carbon isotope incorporation rate across all the intermediates in *E. diaphana* (84.53% ± 1.15%, *p* = 0.0046) and *C. andromeda* (91.20% ±1.29%, *p* = 0.00039), while there was no further increase observed for *S. pistillata* (84.43% ± 2.97%, *p* = 0.45). In addition, significant increases were also observed for ^15^N incorporation rates. In particular, most of the ^15^N isotope was identified in both ^13^C- and ^15^N-containing metabolites (^13^C^15^N) from animals with the combined treatment, while only a small proportion of the ^15^N isotope ended up in ^12^C^15^N compounds (**Tables S1**-**S4**), which indicates that most of the ^15^N was assimilated through the incorporation into carbon backbones derived from the ^13^C_6_-glucose provided. This finding further supported the hypothesis that the metabolization of glucose to 3-phosphohydroxypyruvate produces the carbon backbones required for ammonium assimilation through the GS/GOGAT cycle.

To further determine if the observed assimilation of ammonium is driven by the host animals, we examined the incorporation of ^13^C and ^15^N isotopes in aposymbiotic *E. diaphana* following the same experimental design (**Figure 3B**). Aligned with the patterns observed in symbiotic animals, we found that 92.65% ± 0.79% of the isolated 3-phosphohydroxypyruvate contained ^13^C isotope in aposymbiotic sea anemones. This proved that the ^13^C isotope was integrated directly through the uptake and consumption of ^13^C_6_-glucose by the host. Moreover, the downstream intermediate metabolites showed a significant proportion of ^15^N-containing compounds (^12^C^15^N and ^13^C^15^N), especially in the ^13^C^15^N form (**Figure 3B, Tables S1-S4**). This provided further proof that the animal hosts are able to assimilate the provided ammonium using carbon backbones derived from glucose metabolism, as hypothesized. The similar patterns observed between symbiotic and aposymbiotic sea anemones provide additional proof that the incorporation of ammonium into amino acids is driven by the host animals and that this process does not require the presence of symbionts as long as carbon backbones are provided. Although the presence of symbionts increased the co-incorporation of ^13^C and ^15^N in the combined treatment (**Figure 3B, Tabels S1-S4**), the incorporation rates between symbiotic and aposymbiotic anemones were not significantly different (*t* = -0.40, *p* = 0.69).

### Metabolic destinations for glucose taken up by corals

Besides validating our hypothesis of host-driven ammonium assimilation as a universal mechanism in cnidarian symbioses, we also noticed that corals appear to differ in their use of glucose compared to sea anemones and jellyfish. To track the metabolic flux of ^13^C in our target GS/GOGAT pathway, we calculated the proportional changes relative to total carbon atoms across the pathway metabolites (**Figure 4**). *S. pistillata* showed significantly higher ^13^C uptake and metabolization but significantly lower ^13^C integration into downstream metabolites compared to *E. diaphana* and *C. andromeda* when supplemented with ^13^C_6_-glucose (**Figure 4A**). The relative proportion of ^13^C exhibited a sharper decrease over the whole pathway, as reflected in a significantly steeper slope based on a generalized linear model (**Figure S26**; *S. pistillata* vs *E. diaphana*, -0.112 vs -0.047, *p* = 0.024; *S. pistillata* vs *C. andromeda*, -0.112 vs -0.062, *p* = 0.043). These suggested that corals use relatively more glucose for purposes other than amino acid biosynthesis. Furthermore, the simultaneous provision of ^15^N-ammonium and ^13^C_6_-glucose did not increase the uptake of ^13^C or its integration into amino acids in *S. pistillata* as observed for *E. diaphana* or *C. andromeda* (**Figures 3B** **and 4B**). Conversely, sea anemones and jellyfish showed a higher relative capacity to integrate ^13^C into amino acids and this capacity was further enhanced when additional ammonium was provided.

**Figure 4.**
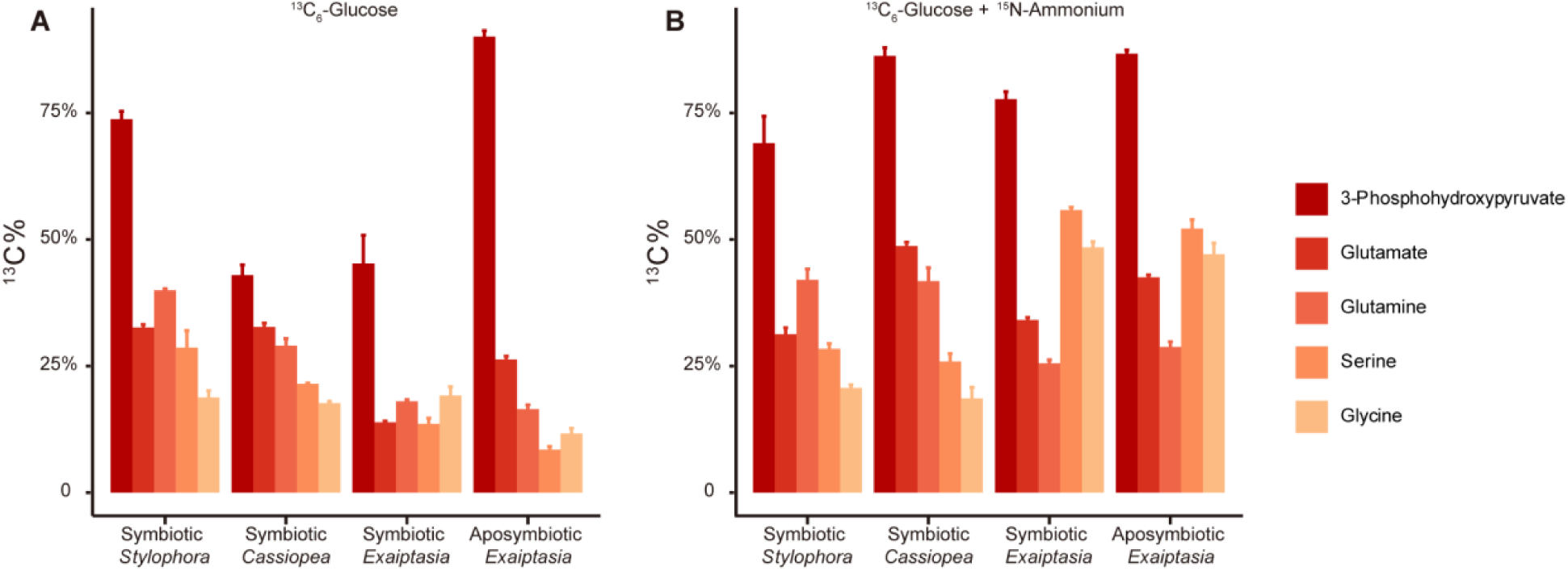
The incorporation of ^13^C across metabolites of GS/GOGAT-mediated amino acid synthesis. Bars represent the standard error of the mean.

## Discussion

Endosymbiotic relationships are the most intimate form of symbiosis, as the symbionts are maintained within the host’s cells (Nowack and Melkonian, 2010). Naturally, this intimacy requires mechanisms that allow providing mutual advantages to both parties in order to maintain an evolutionary stable relationship while discouraging or penalizing parasitic traits that could destabilize the relationship and trigger a Red Queen’s race between host and symbiont (Van Valen, 1973). Oftentimes, this is prevented through the evolution of mechanisms that provide the host with means to control or limit symbiont proliferation in order to prevent the overproliferation of symbionts at the host’s expense (Bull and Rice, 1991; Sachs et al., 2010). Here, we tested our hypothesis that the widespread symbiotic relationships between cnidarians and their dinoflagellate symbionts in the family Symbiodiniaceae are based on a simple metabolic model. This basic metabolic interaction allows cnidarian hosts to control symbiont proliferation while at the same time maximizing their capacity to assimilate and recycle scarce nitrogen into valuable amino acids. This metabolic interaction also provides a parsimonious explanation for the repeated evolution of the symbiotic associations across many cnidarian taxa, and potentially also other marine invertebrates.

While our results clearly show that all four hosts employ ammonium assimilation as a means to control symbiont proliferation, they also revealed important differences between the different cnidarian hosts and their use of glucose and ammonium. These differences have critical implications for the ability of the hosts to control their symbiont populations and, thus, stabilize the symbiosis. Specifically, we found that the rates at which these metabolites are taken up and metabolized differ substantially between the taxa studied, with the coral *S. pistillata* showing the highest uptake of ^13^C_6_-glucose but the lowest relative incorporation rate of ^13^C into ^15^N-containing metabolites. This difference in the incorporation of ^13^C likely results from physiological differences in carbon requirements and utilization. Reef building corals, like *S. pistillata*, require a significant amount of glucose to meet the energy demands of the calcification processes. The significantly lower relative incorporation of ^13^C in more downstream pathway metabolites in comparison to anemones and jellyfish indeed suggests that corals use a larger part of the glucose, and the derived carbon backbones, to meet their energetic demands. Conversely, jellyfish and sea anemones do not calcify and do thus not have such a physiological demand. Hence, they’re able to use more of the glucose provided for the assimilation of ammonium and subsequent amino acid biosynthesis. In the glucose treatment without additional ammonium, these non-calcifying species are rather nitrogen-limited as the provided glucose is sufficient to cover both the energetic demands as well as the assimilation of the ammonium available. However, the provision of both glucose and ammonium further promoted the uptake of both nutrients from the surroundings and subsequent ammonium assimilation and amino acid biosynthesis i.e. more glucose was taken up if additional ammonium was provided, which further increased ammonium assimilation and amino acid biosynthesis.

Our results, therefore, imply that coral hosts might have a lower capacity to assimilate ammonium as they require more glucose to meet their energetic demands, which translates in a lower capacity to control nitrogen levels and, thus, symbiont proliferation. This lowered capacity could also result in a reduced ability to buffer imbalances in the availability of glucose and ammonium. This means that the symbiotic relationship between corals and Symbiodiniaceae is likely more sensitive to metabolic imbalances, i.e. changes in the fluxes of glucose and ammonium, which can result from reduced translocation of photosynthates during heat stress (Rädecker et al., 2021; Tremblay et al., 2016). Interestingly, corals (*S. pistillata* and *A. hemprichii*) consistently showed the strongest response to glucose provision with a dramatic decrease in symbiont density. This fast response suggests that corals might possess a mechanism to actively reduce symbiont density when alternative carbon sources are available. Such a mechanism might allow corals to mitigate some of the drawbacks resulting from their reduced ability to control their symbiont population. In line with this, a recent study suggests that corals might also be able to control ammonium fluxes to the symbionts by varying the expression level of an ammonium transporter at the symbiosome membrane (Thies et al., 2022).

Unlike the other endosymbiosis that is driven by the complementation of the host’s limited metabolic capabilities (Douglas, 2009; Hoffmeister and Martin, 2003; Mao et al., 2018), our findings suggest that cnidarian hosts rely rather heavily on the provision of glucose to control their symbiont population. This host-driven mechanism provides an effective metabolic strategy to gain control over the symbiotic relationship at the expense of being dependent on symbiont-derived glucose. This paradoxical interaction in which control over symbiont proliferation requires symbiont derived photosynthates might also explain the general sensitivity of these relationships to environmental stressors that affect symbiont productivity or nutrient balance (Baker et al., 2018).

## Materials and Methods

### Cnidarian animals

Multiple colonies of the coral *Stylophora pistillata* and *Acropora hemprichii* were collected in the central Red Sea (Al Fahal Reef, 22°14’54”N 38°57’46”E). The coral colonies were acclimatized in indoor tanks with constant sediment-filtered Red Sea water in-flow (salinity ∼39-40 ppt) for at least three months before being used in this study.

The sea anemone *Exaiptasia diaphana* (strain CC7) was used in this study. Aposymbiotic *E. diaphana*, sea anemones free of symbionts, were generated following a cold-shock protocol (Cui et al., 2019). Briefly, animals were treated at 4 °C for 4 h, followed by about 30 days of treatment in 50 μM Diuron with daily water changes. Aposymbiotic animals were maintained under 12 h:12 h light:dark cycle (see below) to asure that no residual symbionts were present and analyzed via fluorescence microscopy to further confirm the absence of algal fluorescence before experimentation. All animals, symbiotic and aposymbiotic, were kept at 25 °C on a 12 h:12 h light:dark cycle with ∼40 μmol photons m^-2^s^-1^ of photosynthetically active radiation and fed with freshly hatched brine shrimp, *Artemia*, approximately three times per week. The individuals used in this study were kept in such conditions for at least six months.

The adult jellyfish *Cassiopea andromeda* were collected from the Red Sea (22°20’23.0”N 39°05’31.1”E). The breeding pairs then spawned in the laboratory and fertilized in the autoclaved seawater. The embryos were transferred to a lab incubator at 26 °C and raised until the medusa stage. Different stages of the *C. andromeda*, including polyp, ephyra, and medusa, were raised separately. All animals were maintained in autoclaved seawater at 26 °C on a 12 h:12 h light:dark cycle with ∼40 µmol photons m^-2^s^-1^ of photosynthetically active radiation and fed daily with freshly hatched brine shrimp, *Artemia*.

### Cell density measurement

#### Coral

Cell density changes in response to glucose and ammonium supplementation were measured for two coral species, *S. pistillata* and *A. hemprichii*, respectively. For each species, 8 branches from different coral colonies were cut for each of the three treatments: the ambient seawater control, 10 mM glucose, and 250 µM ammonium chloride (or 10 mM glucose plus 250 µM ammonium chloride to test their combining effects). The coral branches were tied to plastic stands and placed into three transparent Nalgene™ straight-sided wide-mouth polycarbonate jars (Thermo Fisher Scientific). 250 ml seawater from indoor acclimation tanks was used to fill up each of the jars and the water was changed every two days with fresh treatments applied. To ensure efficient gas exchange in these small volumes, we added magnetic stirring bars to the jars before placing them onto a Cimarec™ i Telesystem Multipoint Stirrer (Thermo Fisher Scientific) with constant stirring at 300 rpm. The whole setup was placed in an incubator at 25°C with ∼80 µmol photons m^-2^s^-1^ radiation and a 12 h:12 h light:dark cycle. After 12-days of incubation, coral fragments were airbrushed with a lysis buffer (0.2 M Tris-HCl, pH=7.5; 0.5% Triton-X; 2 M NaCl) to dissociate and lyse the animal tissues. The tissue lysates were sheared by repeated passage through a 25G needle to release the symbionts. 500 µL of each homogenized sample was centrifuged at 8,000*g* for 2 minutes at room temperature.

#### Sea anemones and jellyfish

9 polyps of *E. diaphana* and *C. andromeda* were used for each of the three treatments: the ambient seawater control, 10 mM glucose, and 250 µM ammonium chloride (or 10 mM glucose plus 250 µM ammonium chloride to test their combined effects). The incubation was performed in 6-well plates. 8 ml autoclaved seawater with appropriate treatments was used and refreshed every two days. After 12-days of incubation, animal polyps were homogenized with the above-mentioned lysis buffer using a cordless motor mixer (Thermo Fisher Scientific). The tissue homogenates were sheared by repeated passage through a 25G needle to release the symbionts. 500 µL of each homogenized sample was centrifuged at 8,000*g* for 2 minutes at room temperature.

#### Cell counting

For all the cnidarian animals, symbiont cells in the pellets were counted using a BD LSRFortessa^TM^ Cell Analyzer (BD Biosciences) based on their chlorophyll fluorescence and forward-scatter signals. Host proteins in the supernatants were quantified using a Pierce Micro BCA^TM^ Protein Assay Kit (Thermo Fisher Scientific) according to the manufacturer’s recommendations. Cell density was determined by normalizing the total cell number to total host protein content. The normality of cell density data was tested using the Shapiro-Wilk test followed by Levene’s test of homogeneity of variance. Statistical differences among conditions were calculated using unpaired two-tailed *t* test.

### Isotope labeling and metabolite extraction

To track the uptake and incorporation of ^13^C and ^15^N isotopes, *S. pistillata* fragments, symbiotic and aposymbiotic *E. diaphana* polyps, and *C. andromeda* at the medusa stage were incubated for 48 hours with either filtered seawater, filtered seawater with 10 mM ^13^C_6_-glucose, filtered seawater with 250 µM ^15^N-ammonium, or filtered seawater and 10 mM ^13^C_6_-glucose and 250 µM ^15^N-ammonium. After 48-hours of incubation, the animal tissues were homogenized following the same procedure mentioned above. The homogenates were centrifuged at 10,000*g* for 5 minutes at 4°C to remove the symbionts. Animal tissue lysates in the supernatants were snap frozen using liquid nitrogen. Animal metabolites were then extracted as previously described (Matthews et al., 2017). Briefly, animal tissue homogenates were further lysed in 5 ml milli-Q water and lyophilized using a freeze dryer (Labconco). The lyophilisates were resuspended in 1 ml pre-chilled (−20 °C) 100% methanol, sonicated for 30 minutes at 4 °C in an ultrasonication bath (Branson), and centrifuged at 3,000g for 30 minutes at 4 °C. The supernatants were collected and stored in -80 °C. The pellets were resuspended in 1 ml 50% methanol (−20 °C) and centrifuged at 3,000g for 30 minutes at 4 °C. The supernatants were then combined with those collected from the previous step. The total extracts were then centrifuged at 16,000g for 15 minutes at 4 °C to remove any potential particulates. The supernatants were dried using a speed vacuum concentrator (Labconco) and stored at -80 °C until further processing.

### Ultra-high-performance liquid chromatography-high resolution mass spectrometry

Amino acid standard solutions were prepared by diluting Amino Acid Standard H (Thermo Fisher Scientific) to the concentrations of 25 μM for all amino acids except *L*-cystine (12.5 μM), followed by a 10-fold dilution with 25 % aqueous methanol. Standard solutions for 3-phosphohydroxypyruvate (Sigma-Aldrich) and O-phospho-*L*-serine (Sigma-Aldrich) were individually prepared at a concentration of 2.5 μM in 25 % aqueous methanol. The solutions for host metabolites were prepared with 200 µL of 25 % aqueous methanol and filtered with a 0.2 µm filter before the UHPLC-HR-MS analysis.

Detections of the amino acids and intermediates (3-phosphohydroxypyruvate, and O-phospho-*L*-serine) were performed on a Dionex Ultimate 3000 UHPLC system coupled with a Q Exactive Plus mass spectrometer (Thermo Fisher Scientific) with a heated-electrospray ionization source. Chromatographic separation of amino acids and intermediates was carried out on an ACQUITY UPLC^®^ BEH Amide column (130Å, 1.7 µm, 2.1 mm × 100 mm, Waters) maintained at 35 °C. The mobile phases A (water/formic acid, 100/0.1, v/v) and B (acetonitrile/formic acid, 100/0.1, v/v) were employed for eluting amino acids at a flow rate of 0.25 mL/min and with the gradient program: 0–8 min, 95 % B to 25 % B; 8–11 min, 25 % B; 11–12 min, 25 % B to 95 % B; 12–15 min, 95 % B. In addition, intermediates were eluted with the gradient program: 0–5 min, 100 % B to 25 % B; 5–8 min, 25 % B; 8–9 min, 25 % B to 100 % B; 9–12 min, 100 % B. The injection volume was 2 μL. Amino acids were detected using a mass spectrometer operated in positive mode with a spray voltage of 3.0 kV, sheath gas flow rate of 35 arbitrary units, auxiliary gas flow rate of 10 arbitrary units, spray capillary temperature of 300 °C, auxiliary gas heater temperature of 325 °C, AGC target of 3e6, and resolution of 280,000. In addition, intermediates were detected using a mass spectrometer operated in negative mode with a spray voltage of 2.5 kV, sheath gas flow rate of 40 arbitrary units, auxiliary gas flow rate of 20 arbitrary units, spray capillary temperature of 325 °C, and auxiliary gas heater temperature of 350 °C. In this work, Xcalibur software was used for the MS data acquisition and analysis. Amino acids and intermediates from animal tissues were identified and assigned based on their accurate mass and matching with the corresponding standards. The normalized peak areas of metabolites with that in the natural form were used for their quantitative comparison.

### ^13^C incorporation

We extracted the percentage of ^13^C for each selected metabolite from the GS/GOGAT-mediated amino acid synthesis pathway and then determined its flux for each sample from the appropriate treatments based on a generalized linear regression model using *glm* function implemented in the R base package. The change of incorporation rates was calculated by averaging the slopes estimated from samples in the same treatment. We then conducted pairwise comparisons using a Wilcoxon rank-sum exact test with Bonferroni correction for *p*-value adjustment.

## Author contributions

M.A. and G.C. conceived the study. A.M. and G.C. performed the nutrient supplementation experiments and the algal cell density measurements. G.C. and A.M. performed the isotope-labeling experiments and extracted the metabolites. J.M. and S.A.-B. performed UHPLC-HR-MS experiments. G.C., J.M., A.M., and H.Z. analyzed the metabolomic data. S.H.H. raised the jellyfish line. G.C. and M.A. wrote the initial draft of the manuscript with input from all of the authors. All authors reviewed and edited the manuscript.

## Acknowledgment

Figure 1 was created by Heno Hwang, scientific illustrator at King Abdullah University of Science and Technology (KAUST).

## Competing interests

The authors declare no competing interests.

## Supplemental information

### Supplemental Figures

**Figure S1.**
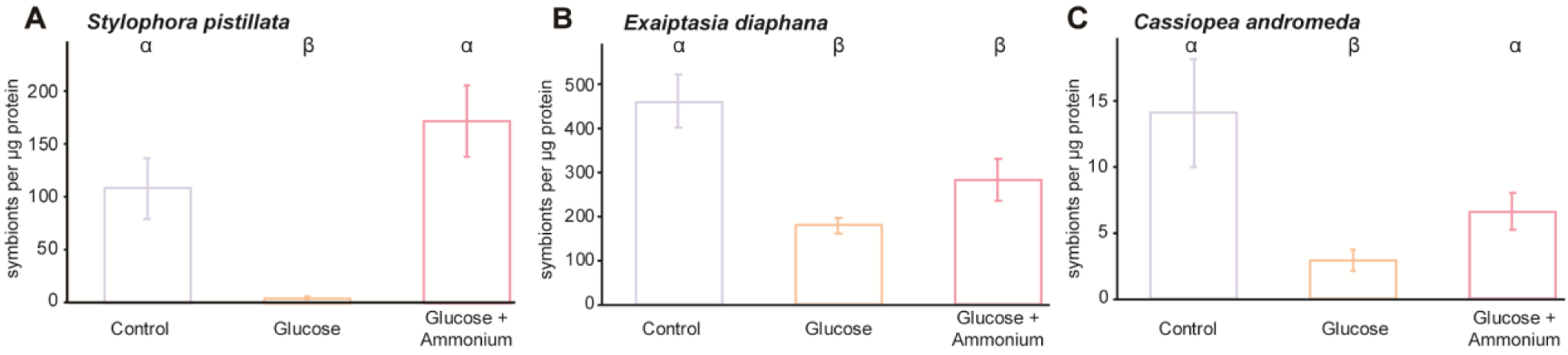
The combined effects of glucose and ammonium on symbiont cell density changes. Symbiont cell densities were calculated by normalizing total symbiont numbers to host protein content. Bars represent the mean ± SE. Greek letters indicate statistical differences with a significance cut-off at *p* = 0.05.

**Figure S2.**
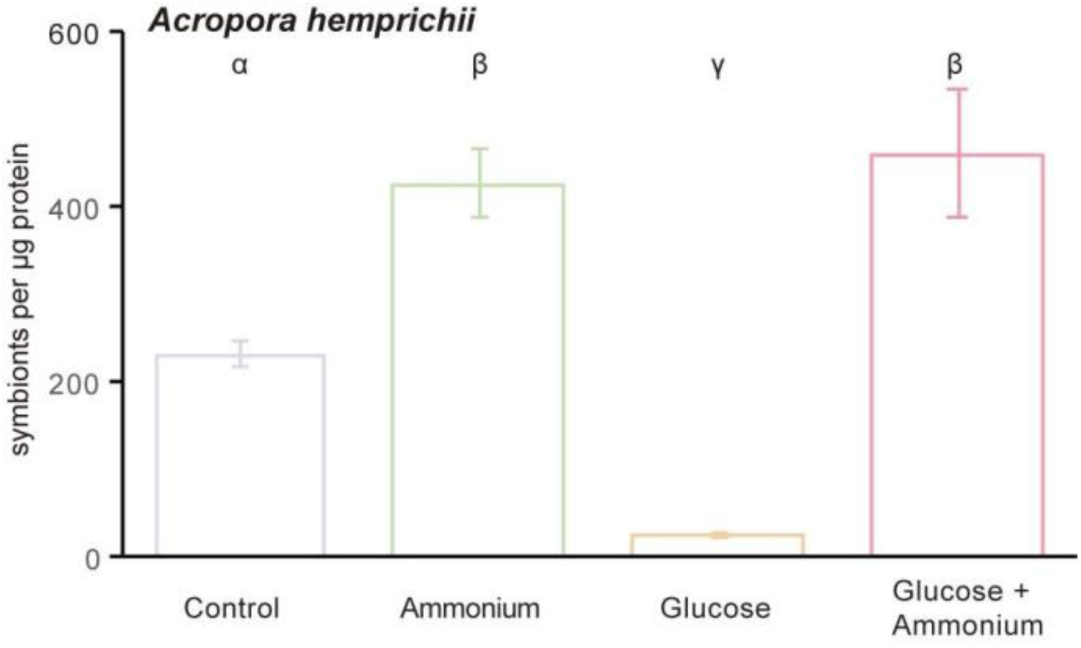
Effects of glucose and ammonium on symbiont cell density changes in the coral *Acropora hemprichii*. Bars represent the mean ± SE. Greek letters indicate statistical differences with a significance cut-off at *p* = 0.05.

**Figure S3.**
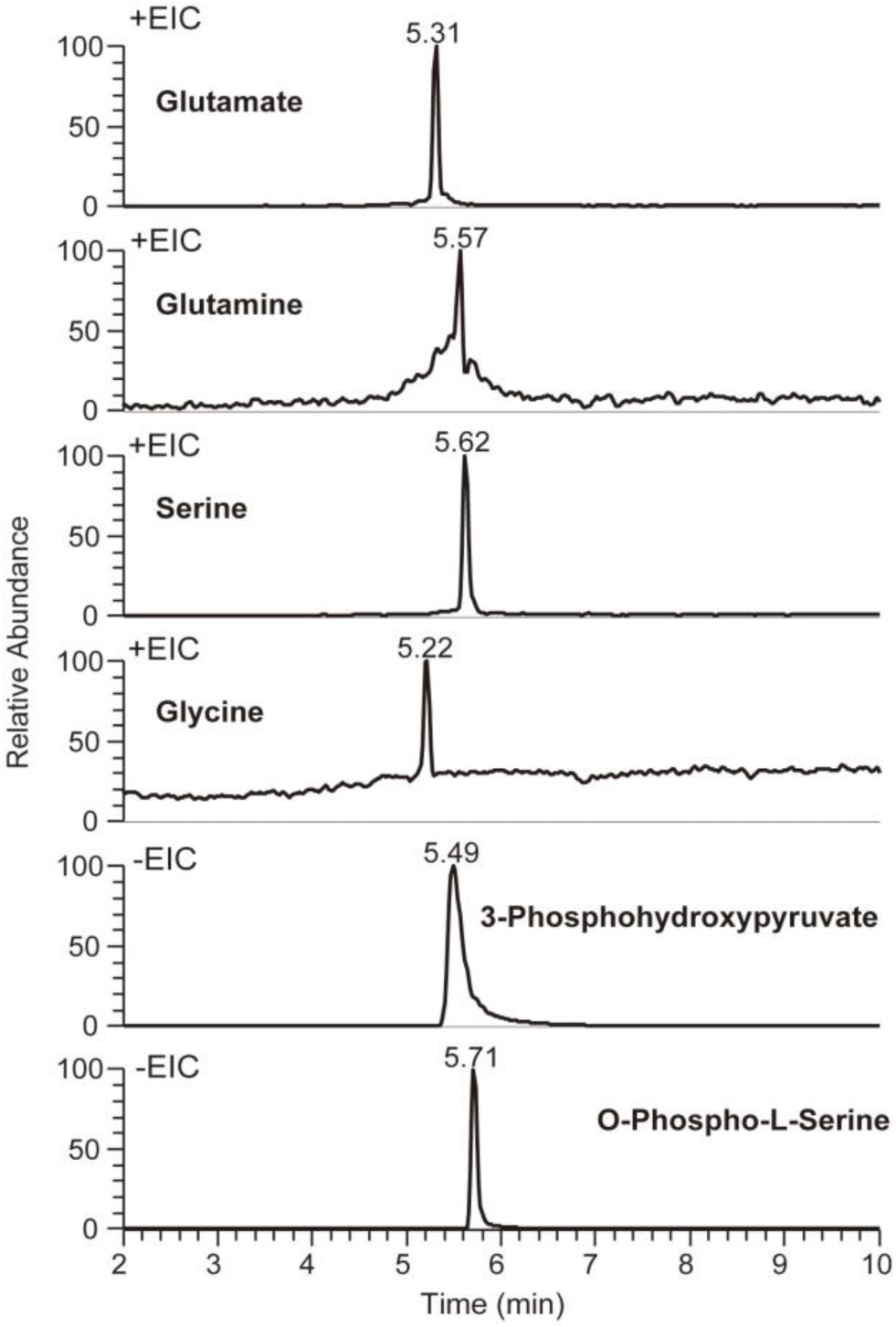
Extracted ion chromatograms (EIC) of standards.

**Figure S4.**
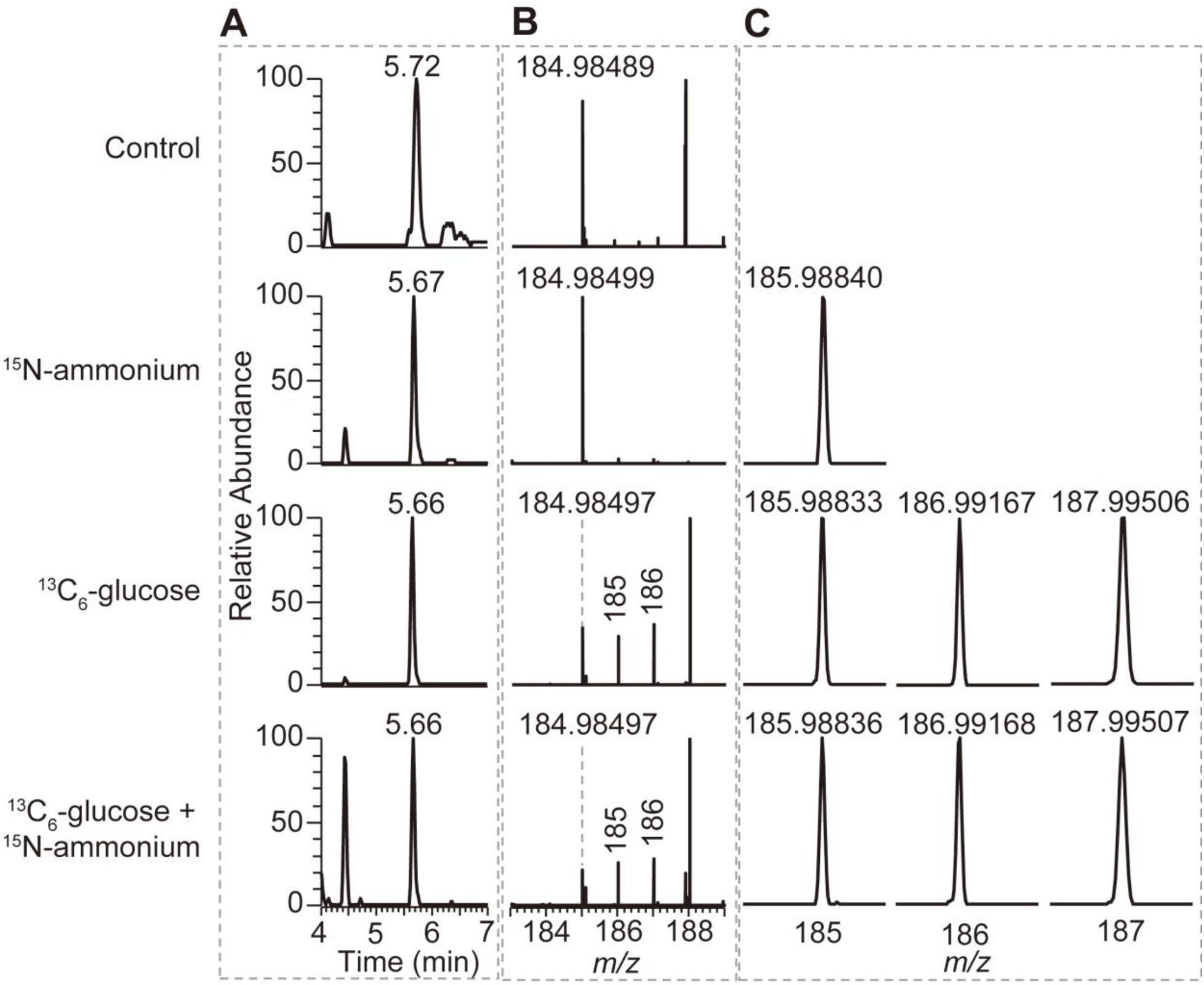
Identification of 3-phosphohydroxypyruvate isolated from symbiotic *S. pistillata*. (**A, B**) Extracted ion chromatograms (EIC) (**A**) and the isotopic distributions (**B**) of 3-phosphohydroxypyruvate isolated from symbiotic *S. pistillata* at different conditions. (**C**) The zoom of the area in (B) showing that different isotopologue compositions of 3-phosphohydroxypyruvate are distinguished by HR-MS.

**Figure S5.**
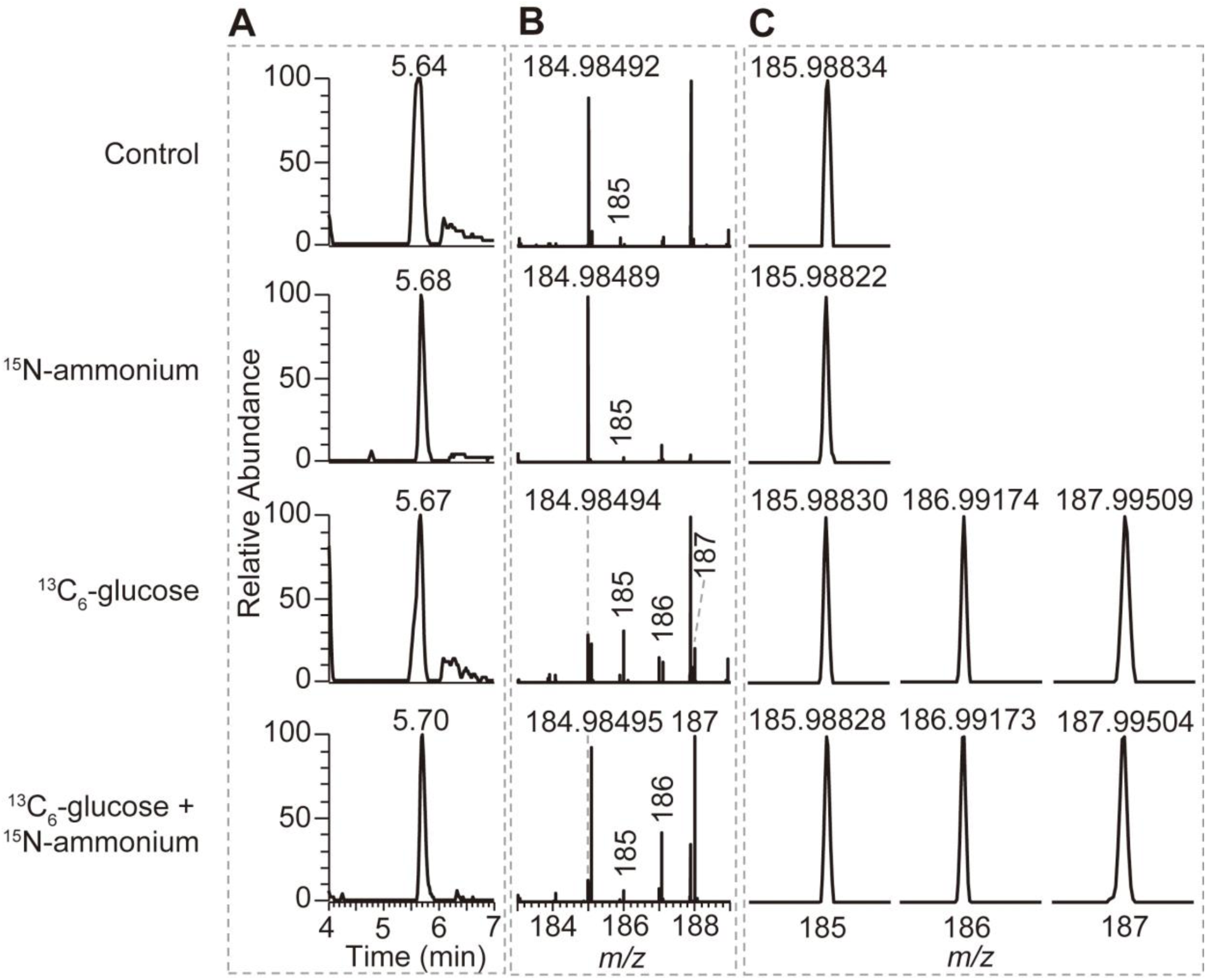
Identification of 3-phosphohydroxypyruvate isolated from symbiotic *C. andromeda*. (**A, B**) Extracted ion chromatograms (EIC) (**A**) and the isotopic distributions (**B**) of 3-phosphohydroxypyruvate isolated from symbiotic *C. andromeda* at different conditions. (**C**) The zoom of the area in (B) showing that different isotopologue compositions of 3-phosphohydroxypyruvate are distinguished by HR-MS.

**Figure S6.**
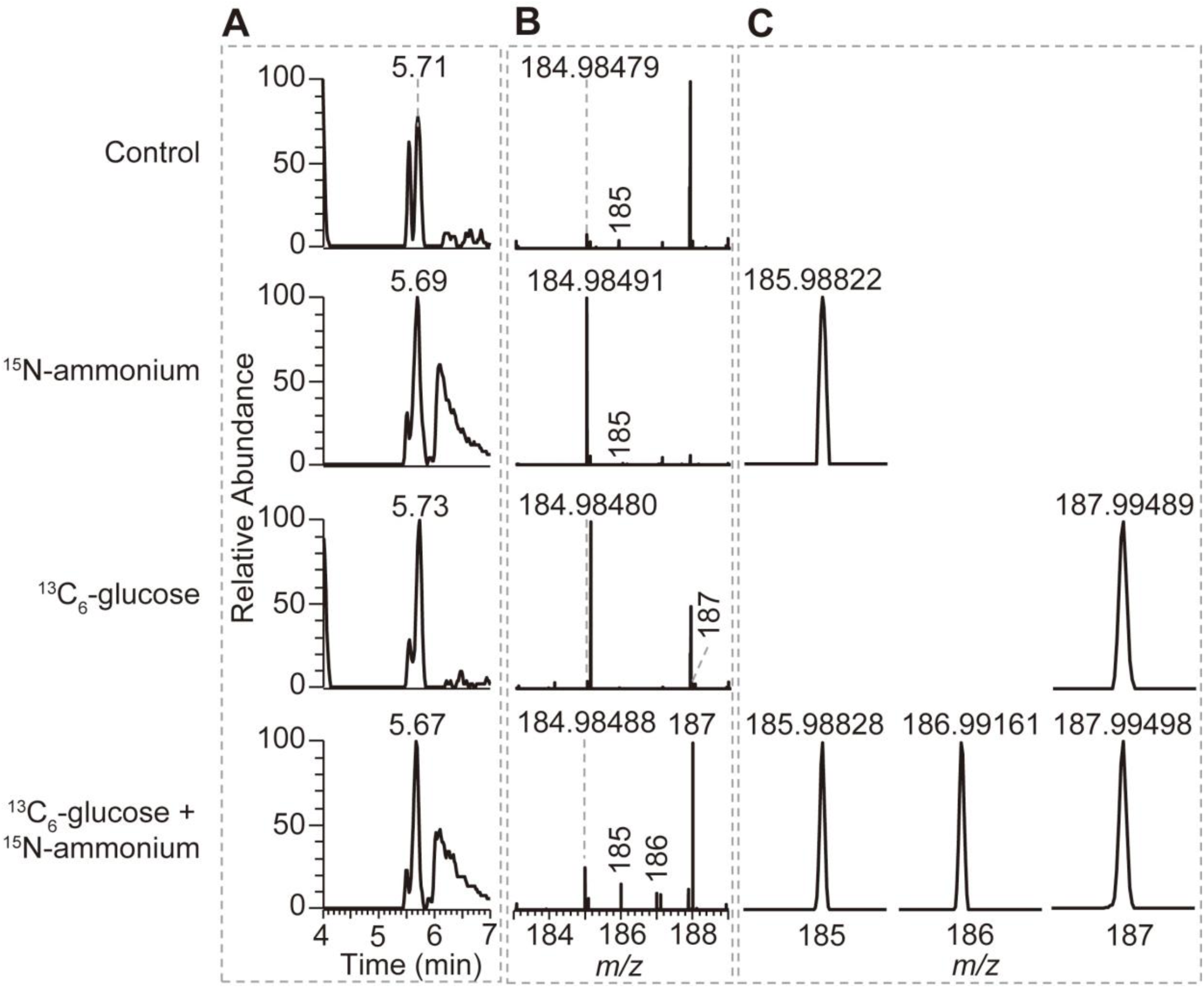
Identification of 3-phosphohydroxypyruvate isolated from symbiotic *E. diaphana*. (**A, B**) Extracted ion chromatograms (EIC) (**A**) and the isotopic distributions (**B**) of 3-phosphohydroxypyruvate isolated from symbiotic *E. diaphana* at different conditions. (**C**) The zoom of the area in (B) showing that different isotopologue compositions of 3-phosphohydroxypyruvate are distinguished by HR-MS.

**Figure S7.**
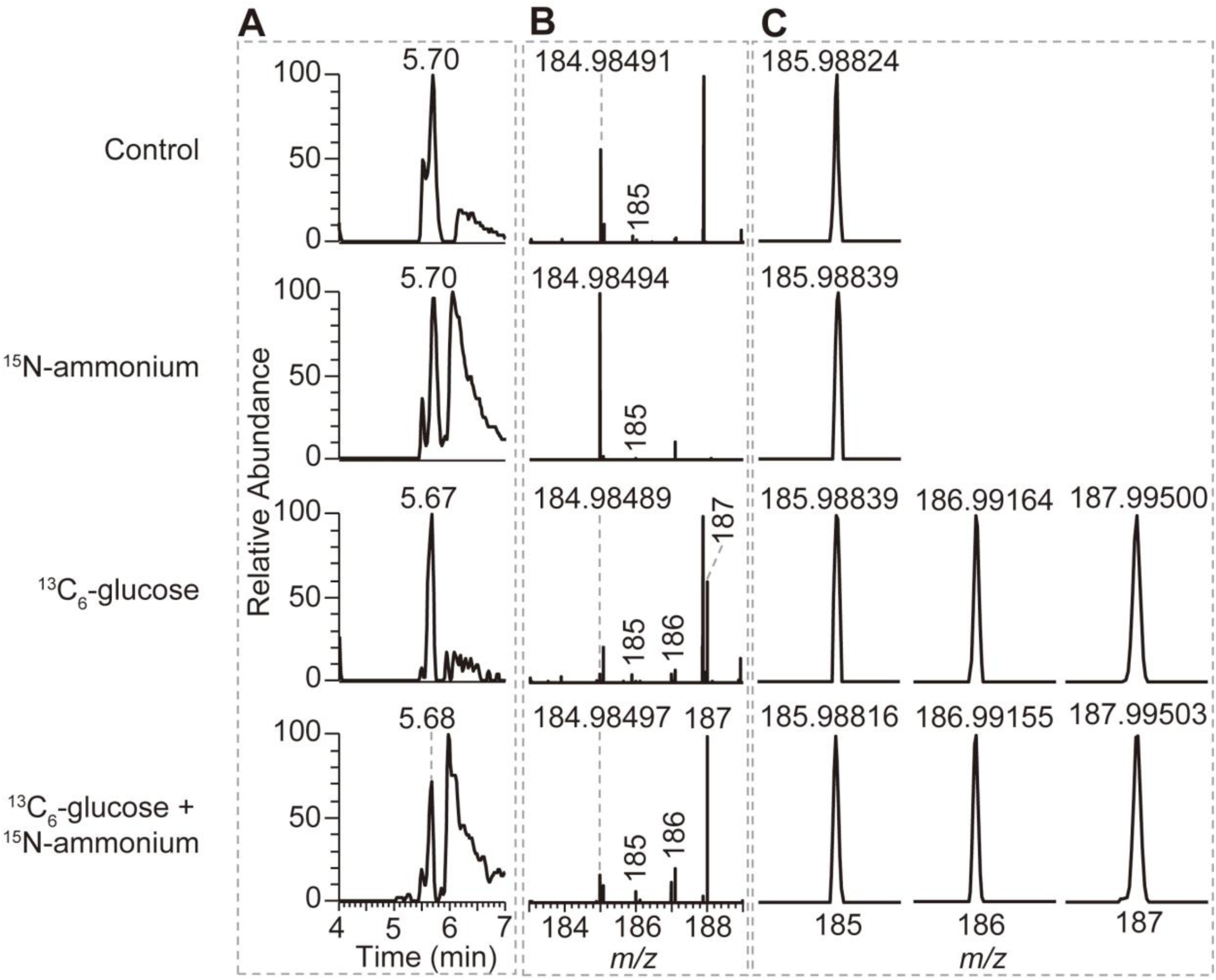
Identification of 3-phosphohydroxypyruvate isolated from aposymbiotic *E. diaphana*. (**A, B**) Extracted ion chromatograms (EIC) (**A**) and the isotopic distributions (**B**) of 3-phosphohydroxypyruvate isolated from aposymbiotic *E. diaphana* at different conditions. (**C**) The zoom of the area in (B) showing that different isotopologue compositions of 3-phosphohydroxypyruvate are distinguished by HR-MS.

**Figure S8.**
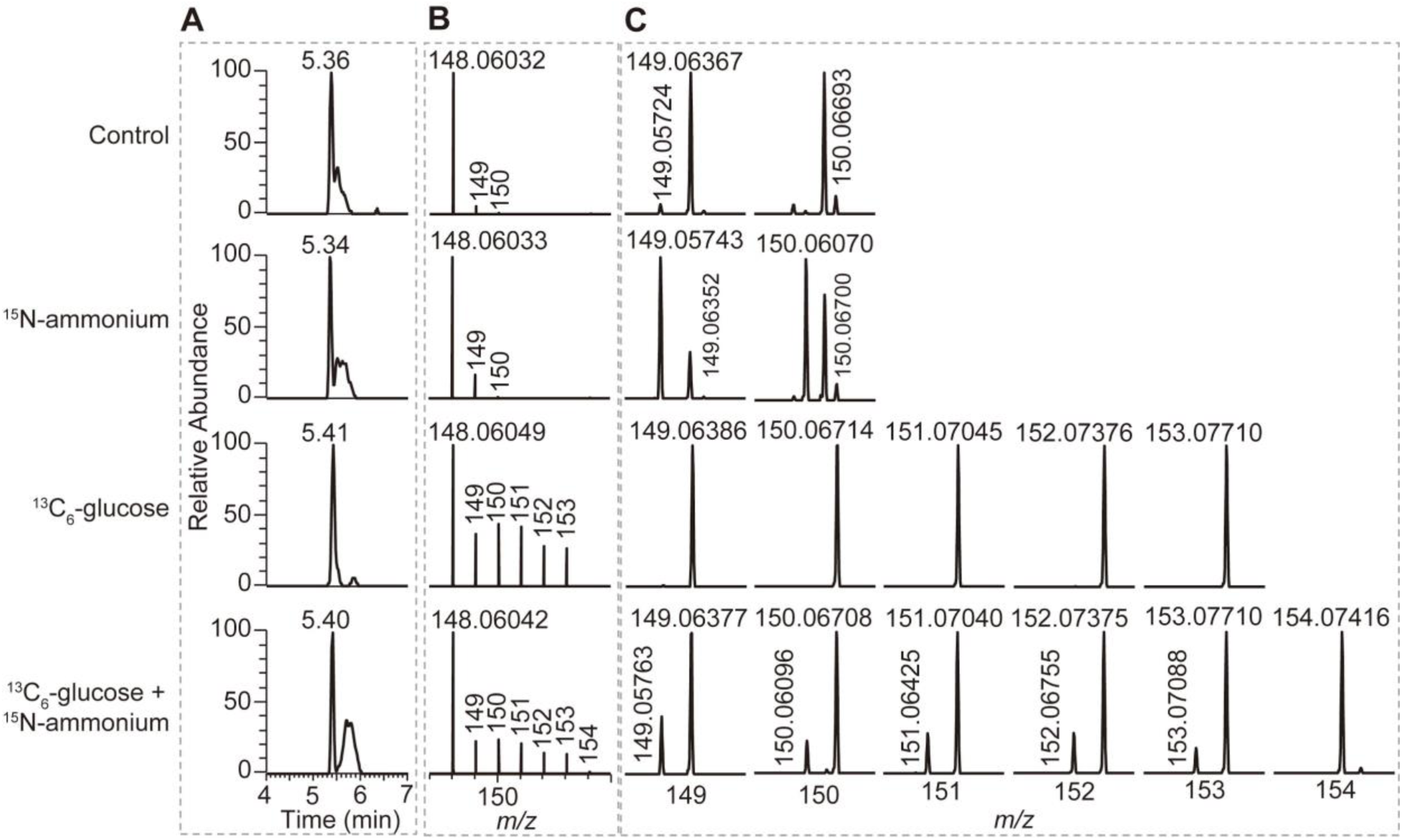
Identification of glutamate isolated from symbiotic *S. pistillata*. (**A, B**) Extracted ion chromatograms (EIC) (**A**) and the isotopic distributions (**B**) of glutamate isolated from symbiotic *S. pistillata* at different conditions. (**C**) The zoom of the area in (B) showing that different isotopologue compositions of glutamate are distinguished by HR-MS.

**Figure S9.**
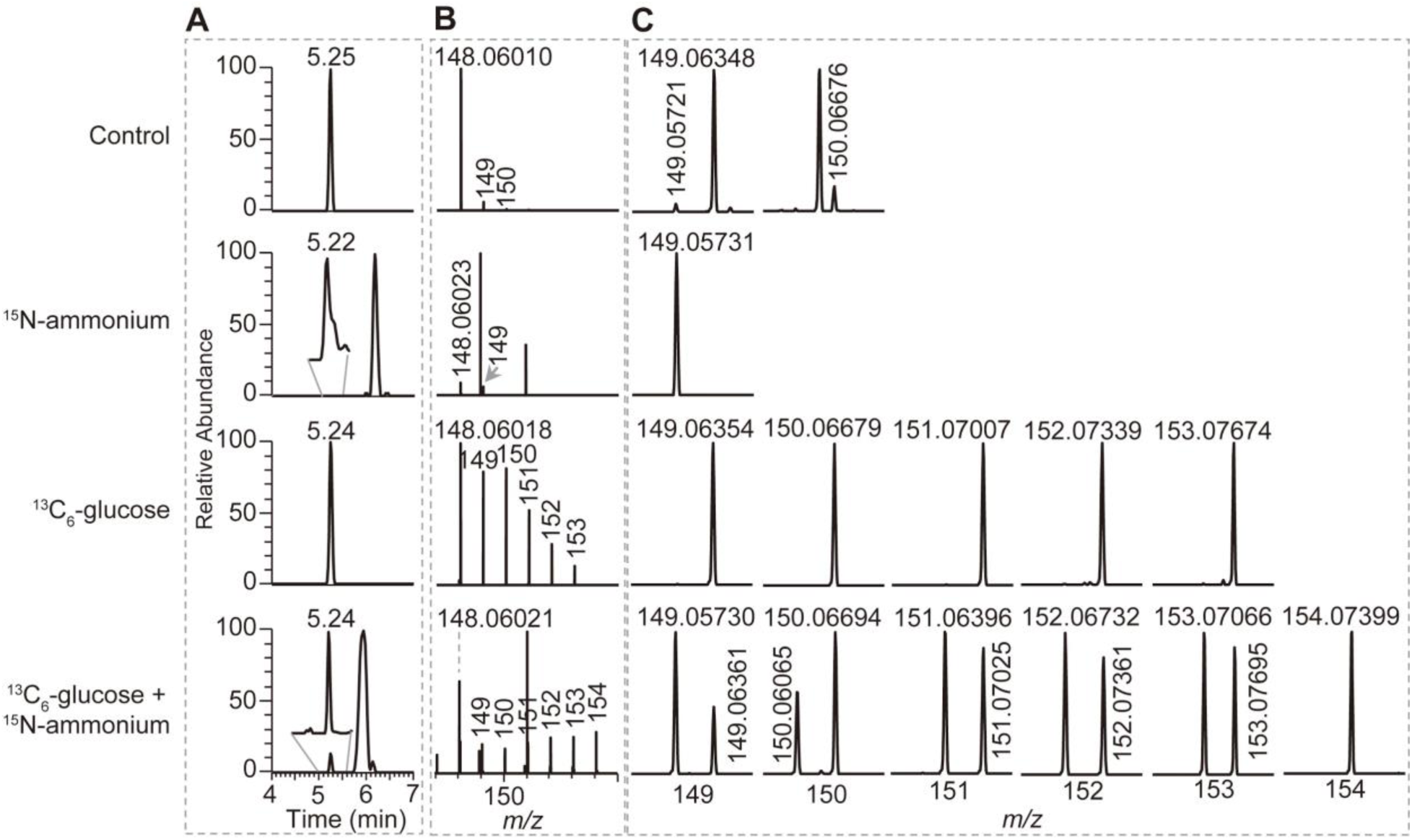
Identification of glutamate isolated from symbiotic *C. andromeda*. (**A, B**) Extracted ion chromatograms (EIC) (**A**) and the isotopic distributions (**B**) of glutamate isolated from symbiotic *C. andromeda* at different conditions. (**C**) The zoom of the area in (B) showing that different isotopologue compositions of glutamate are distinguished by HR-MS.

**Figure S10.**
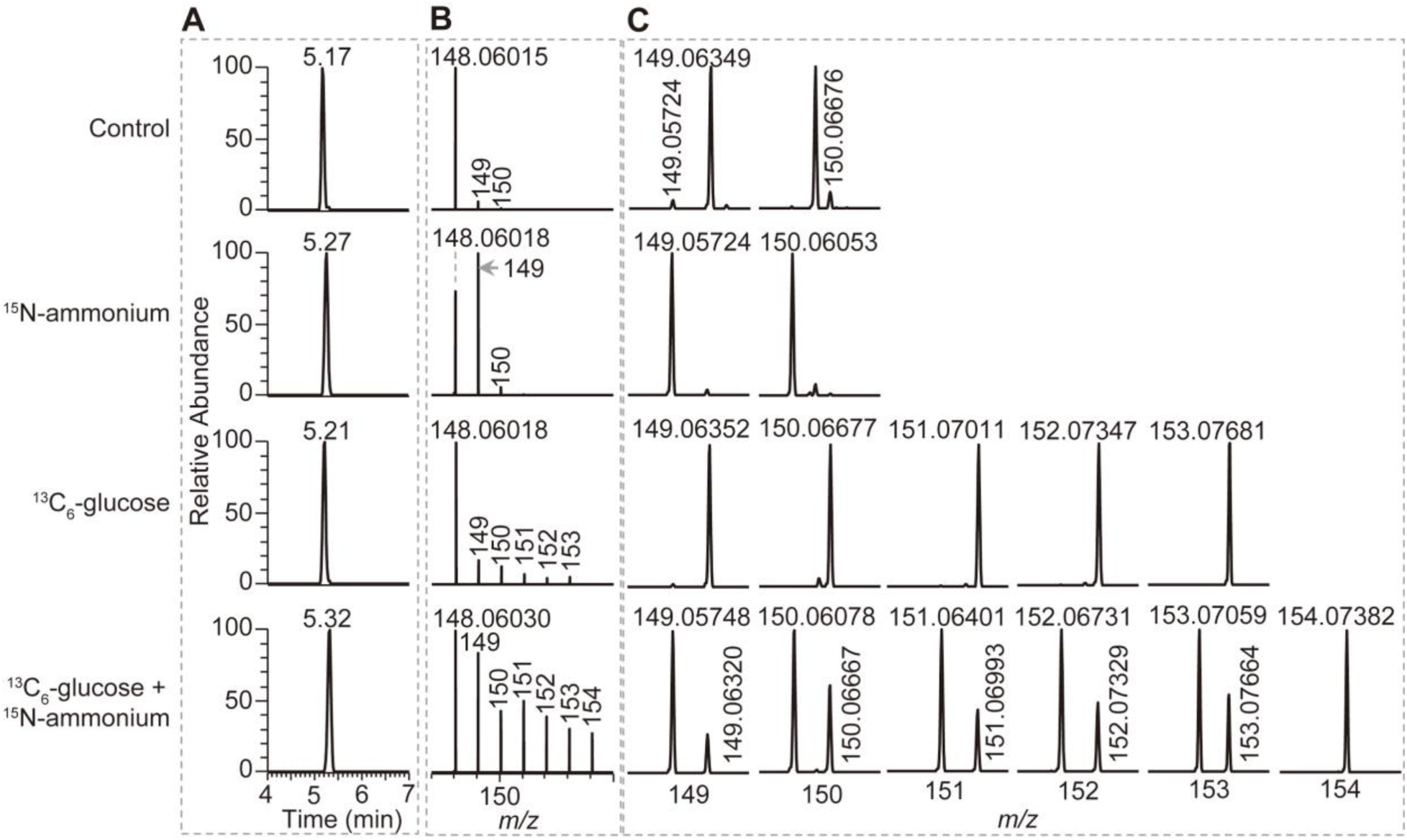
Identification of glutamate isolated from symbiotic *E. diaphana*. (**A, B**) Extracted ion chromatograms (EIC) (**A**) and the isotopic distributions (**B**) of glutamate isolated from symbiotic *E. diaphana* at different conditions. (**C**) The zoom of the area in (B) showing that different isotopologue compositions of glutamate are distinguished by HR-MS.

**Figure S11.**
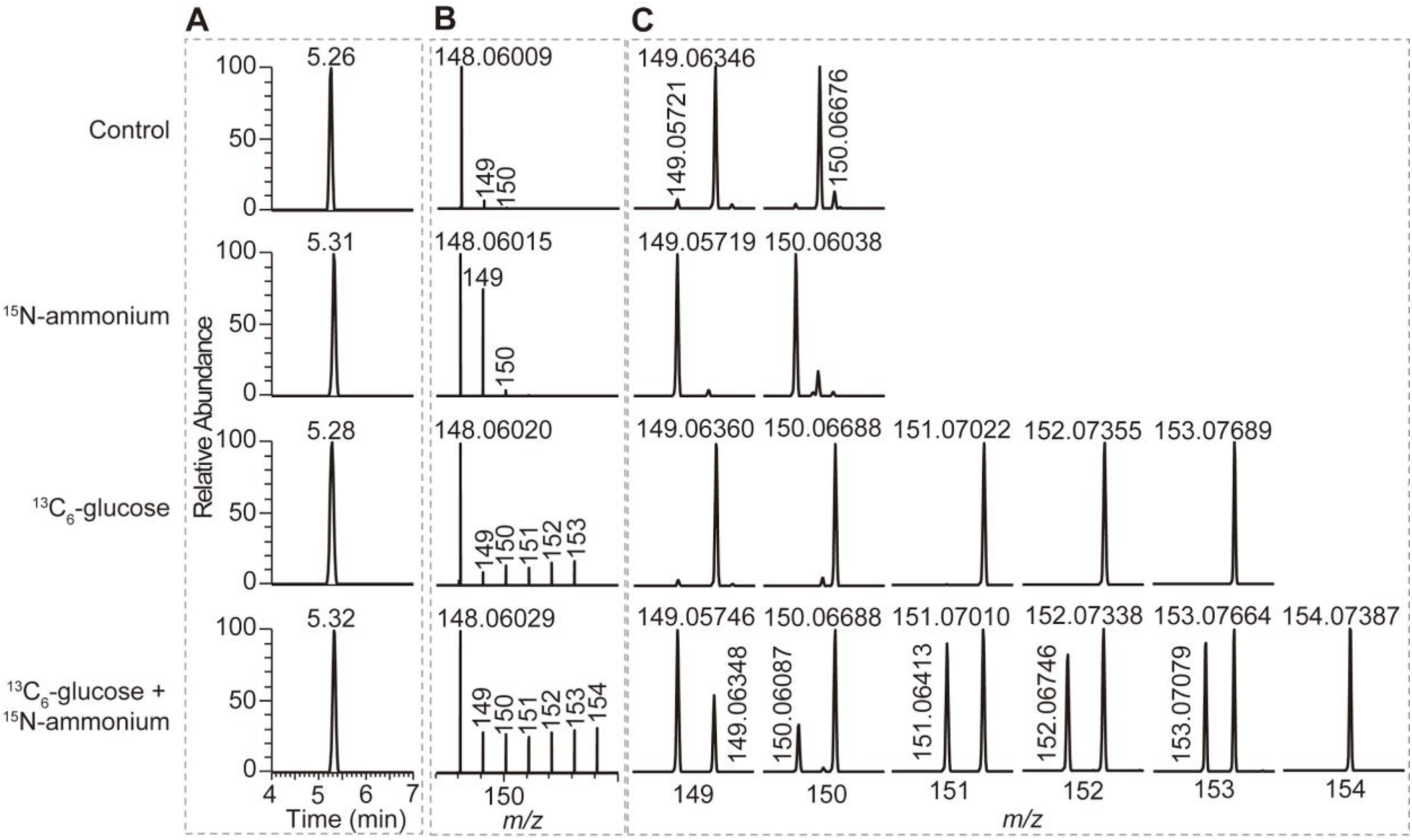
Identification of glutamate isolated from aposymbiotic *E. diaphana*. (**A, B**) Extracted ion chromatograms (EIC) (**A**) and the isotopic distributions (**B**) of glutamate isolated from aposymbiotic *E. diaphana* at different conditions. (**C**) The zoom of the area in (B) showing that different isotopologue compositions of glutamate are distinguished by HR-MS.

**Figure S12.**
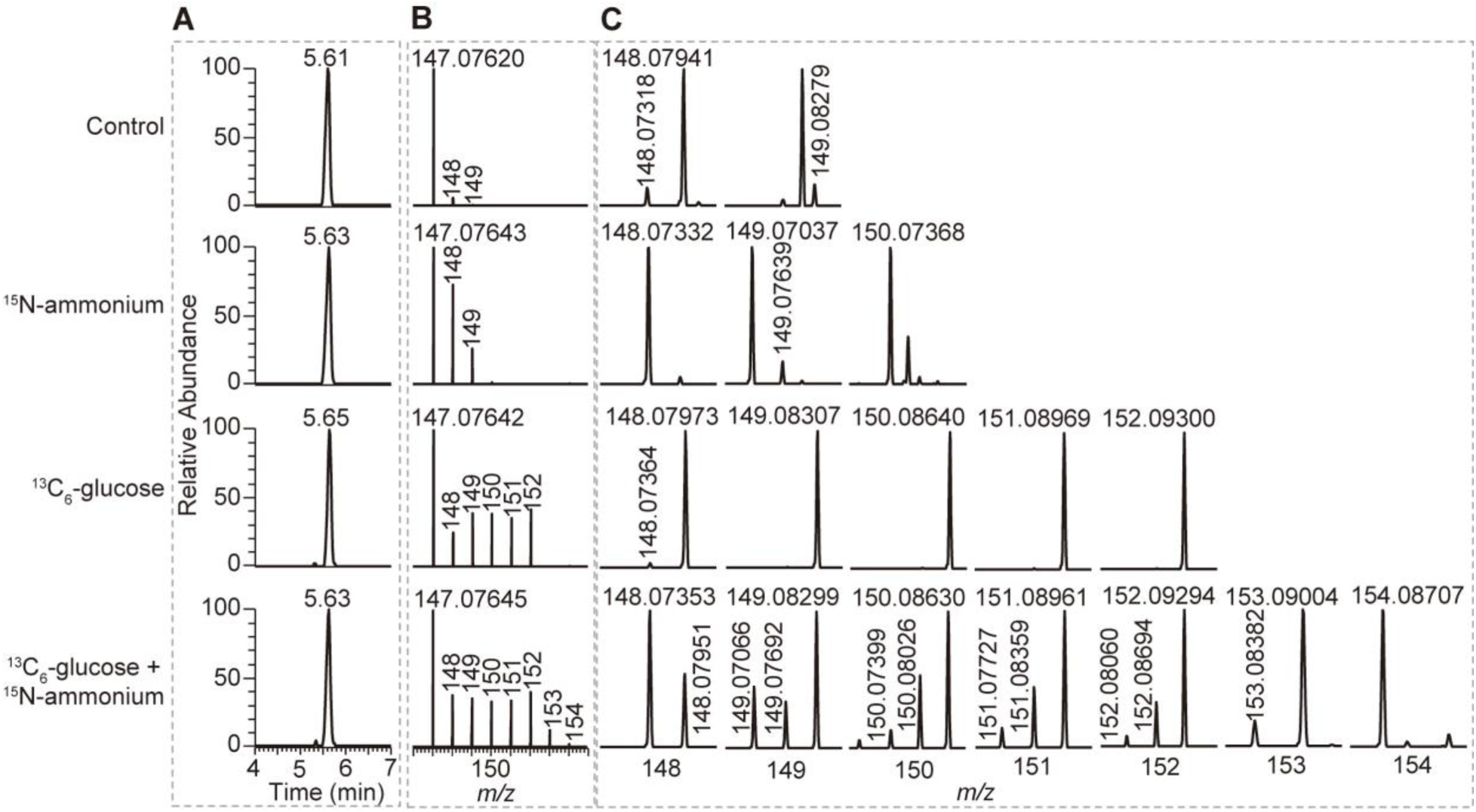
Identification of glutamine isolated from symbiotic *S. pistillata*. (**A, B**) Extracted ion chromatograms (EIC) (**A**) and the isotopic distributions (**B**) of glutamine isolated from symbiotic *S. pistillata* at different conditions. (**C**) The zoom of the area in (B) showing that different isotopologue compositions of glutamine are distinguished by HR-MS.

**Figure S13.**
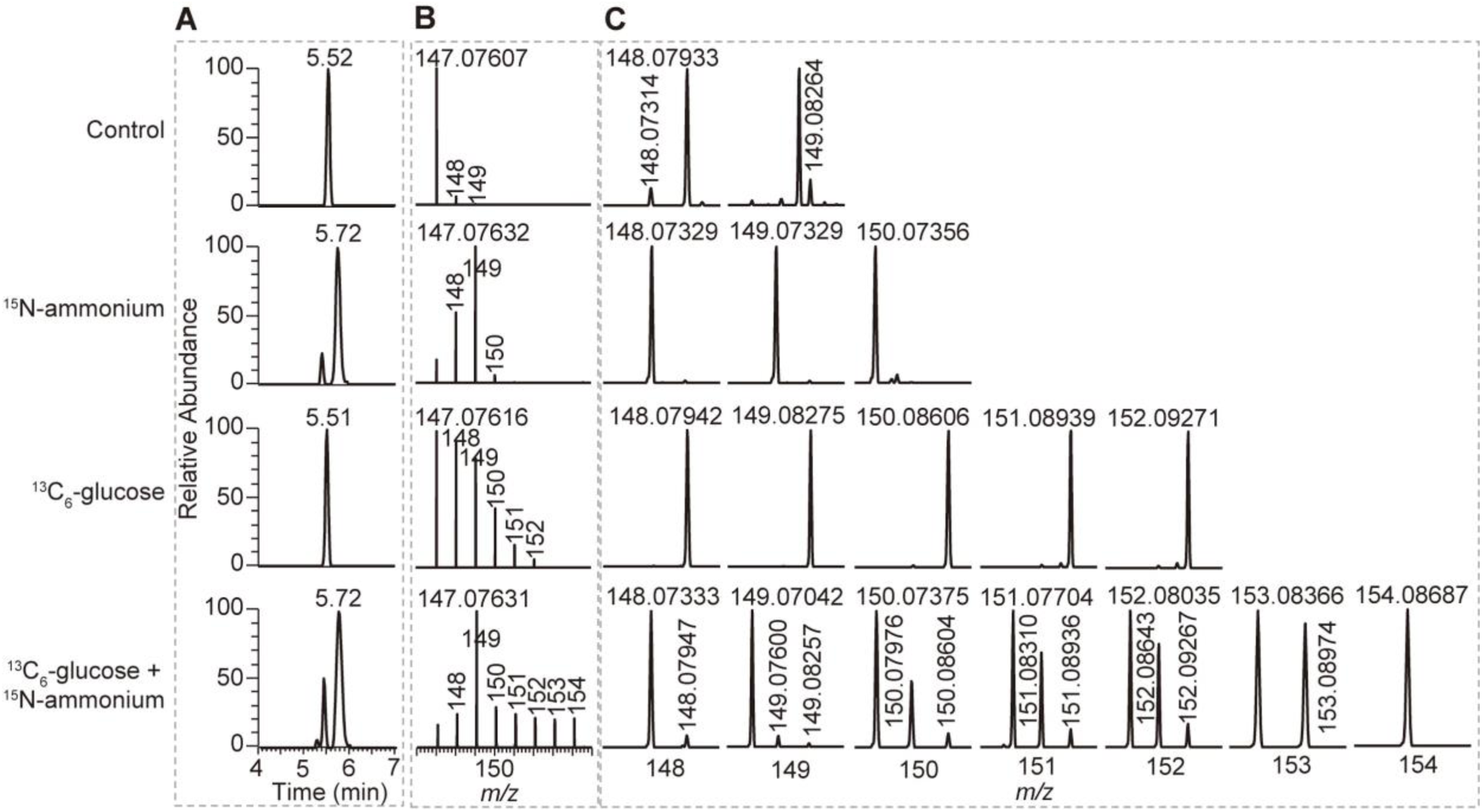
Identification of glutamine isolated from symbiotic *C. andromeda*. (**A, B**) Extracted ion chromatograms (EIC) (**A**) and the isotopic distributions (**B**) of glutamine isolated from symbiotic *C. andromeda* at different conditions. (**C**) The zoom of the area in (B) showing that different isotopologue compositions of glutamine are distinguished by HR-MS.

**Figure S14.**
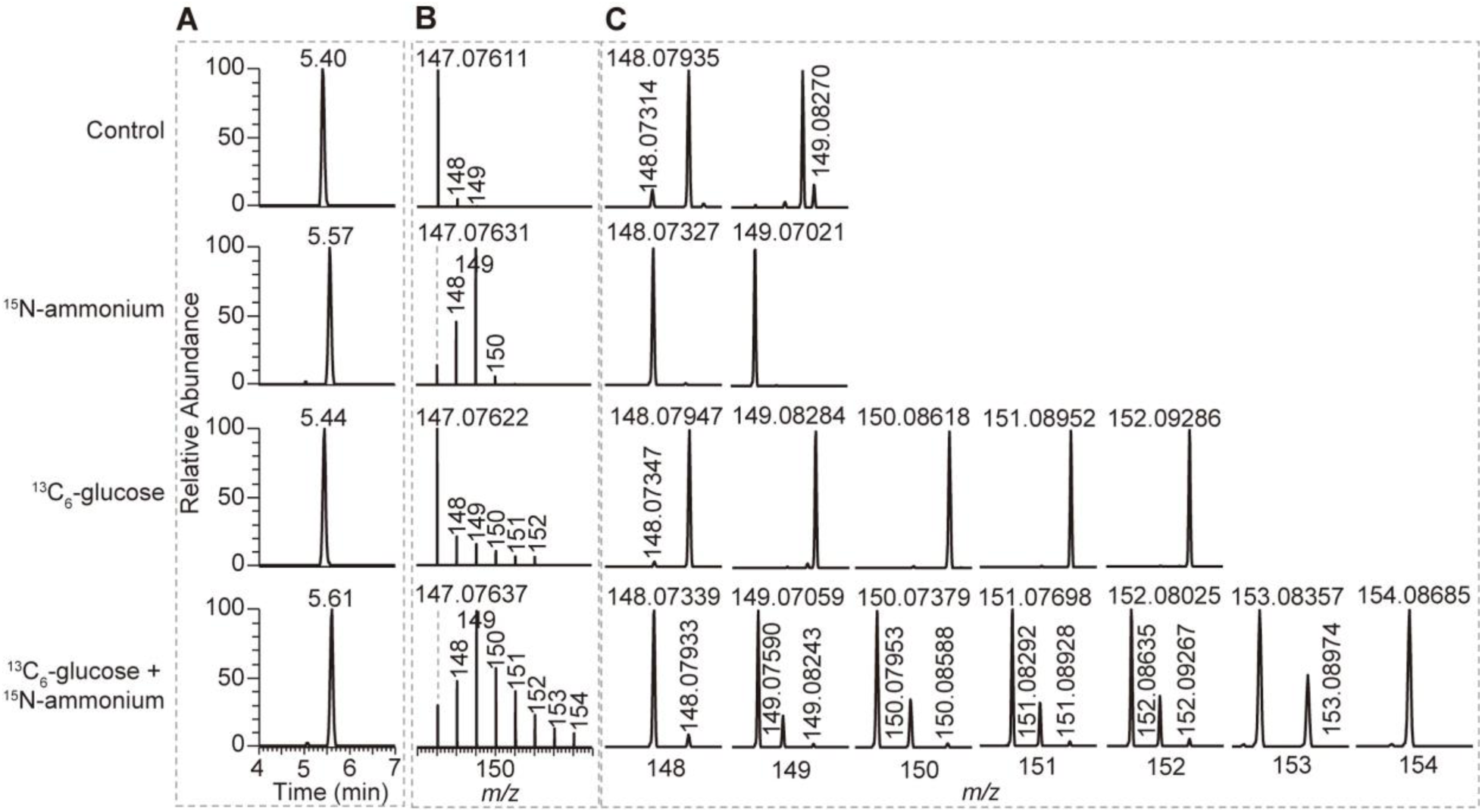
Identification of glutamine isolated from symbiotic *E. diaphana*. (**A, B**) Extracted ion chromatograms (EIC) (**A**) and the isotopic distributions (**B**) of glutamine isolated from symbiotic *E. diaphana* at different conditions. (**C**) The zoom of the area in (B) showing that different isotopologue compositions of glutamine are distinguished by HR-MS.

**Figure S15.**
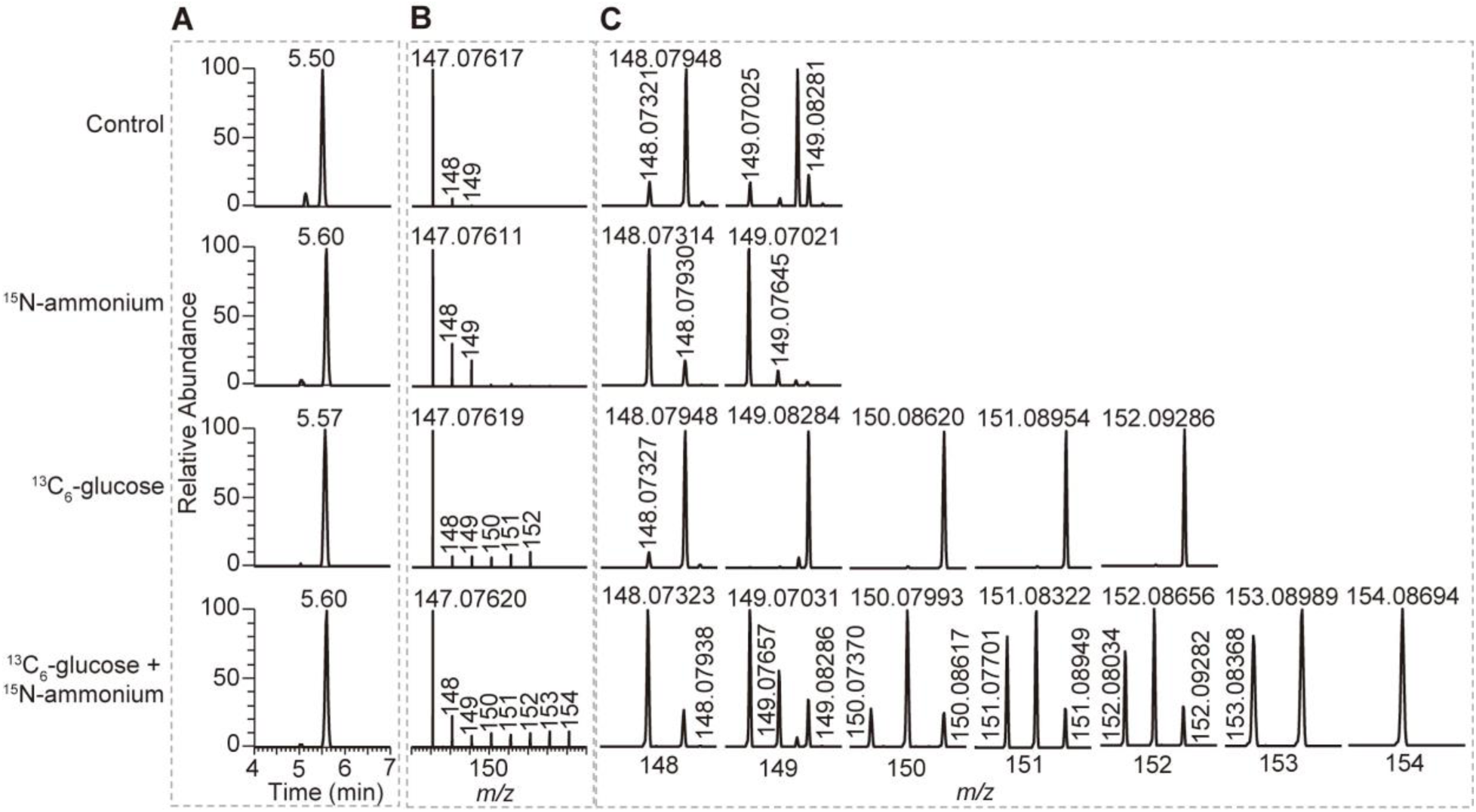
Identification of glutamine isolated from aposymbiotic *E. diaphana*. (**A, B**) Extracted ion chromatograms (EIC) (**A**) and the isotopic distributions (**B**) of glutamine isolated from aposymbiotic *E. diaphana* at different conditions. (**C**) The zoom of the area in (B) showing that different isotopologue compositions of glutamine are distinguished by HR-MS.

**Figure S16.**
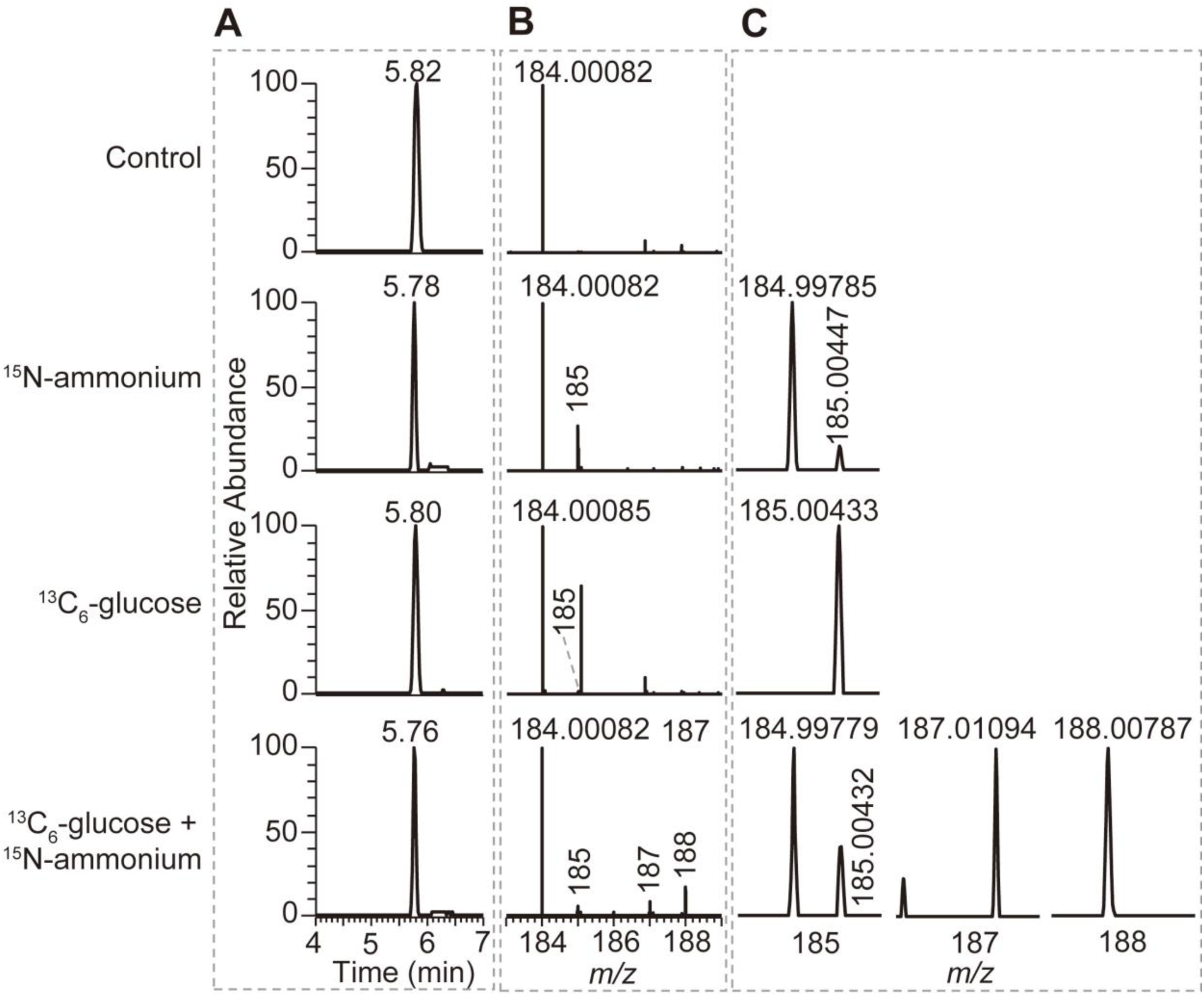
Identification of O-phospho-*L*-serine isolated from symbiotic *E. diaphana*. (**A, B**) Extracted ion chromatograms (EIC) (**A**) and the isotopic distributions (**B**) of O-phospho-*L*-serine isolated from symbiotic *E. diaphana* at different conditions. (**C**) The zoom of the area in (B) showing that different isotopologue compositions of O-phospho-*L*-serine are distinguished by HR-MS.

**Figure S17.**
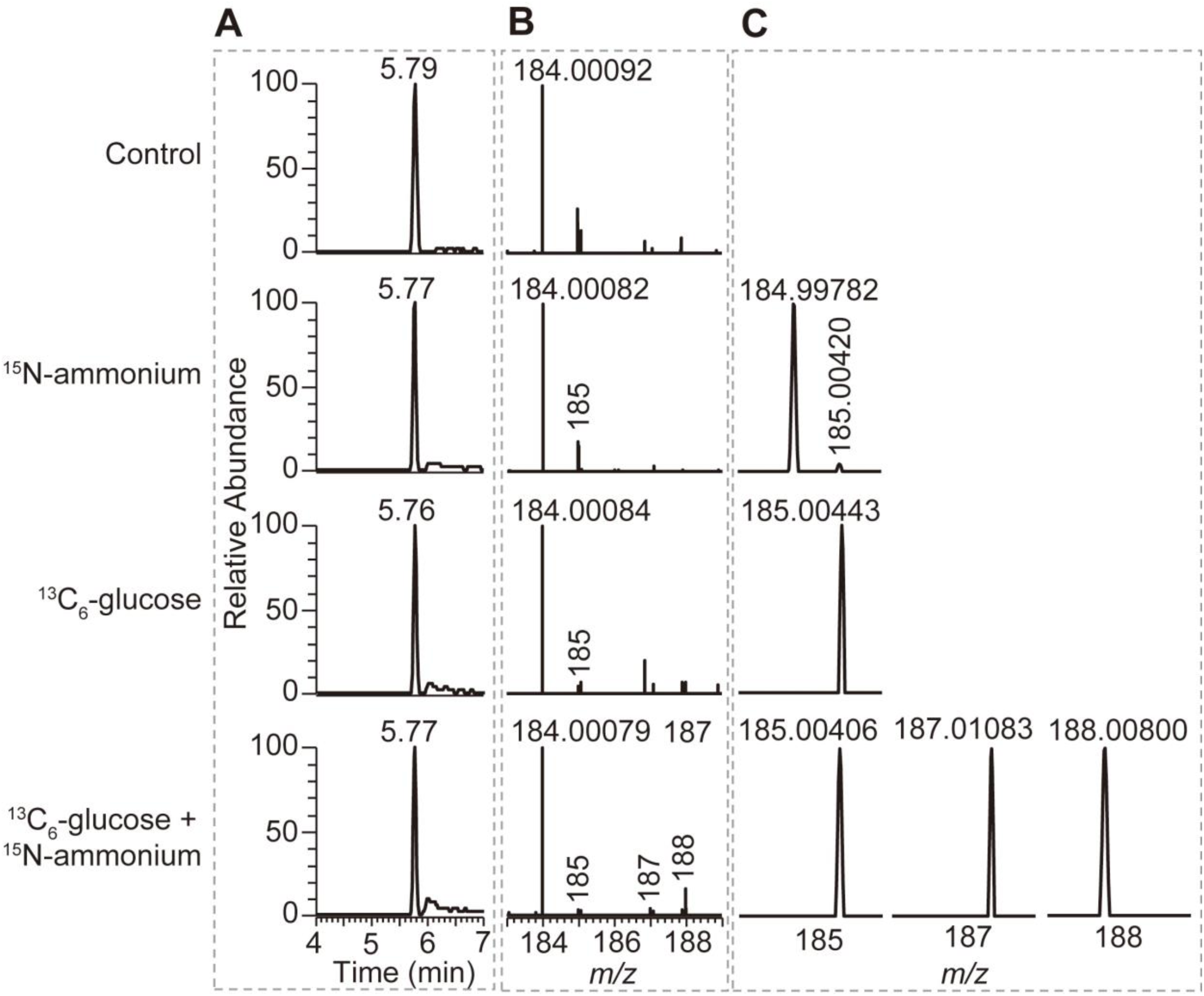
Identification of O-phospho-*L*-serine isolated from aposymbiotic *E. diaphana*. (**A, B**) Extracted ion chromatograms (EIC) (**A**) and the isotopic distributions (**B**) of O-phospho-*L*-serine isolated from aposymbiotic *E. diaphana* at different conditions. (**C**) The zoom of the area in (B) showing that different isotopologue compositions of O-phospho-*L*-serine are distinguished by HR-MS.

**Figure S18.**
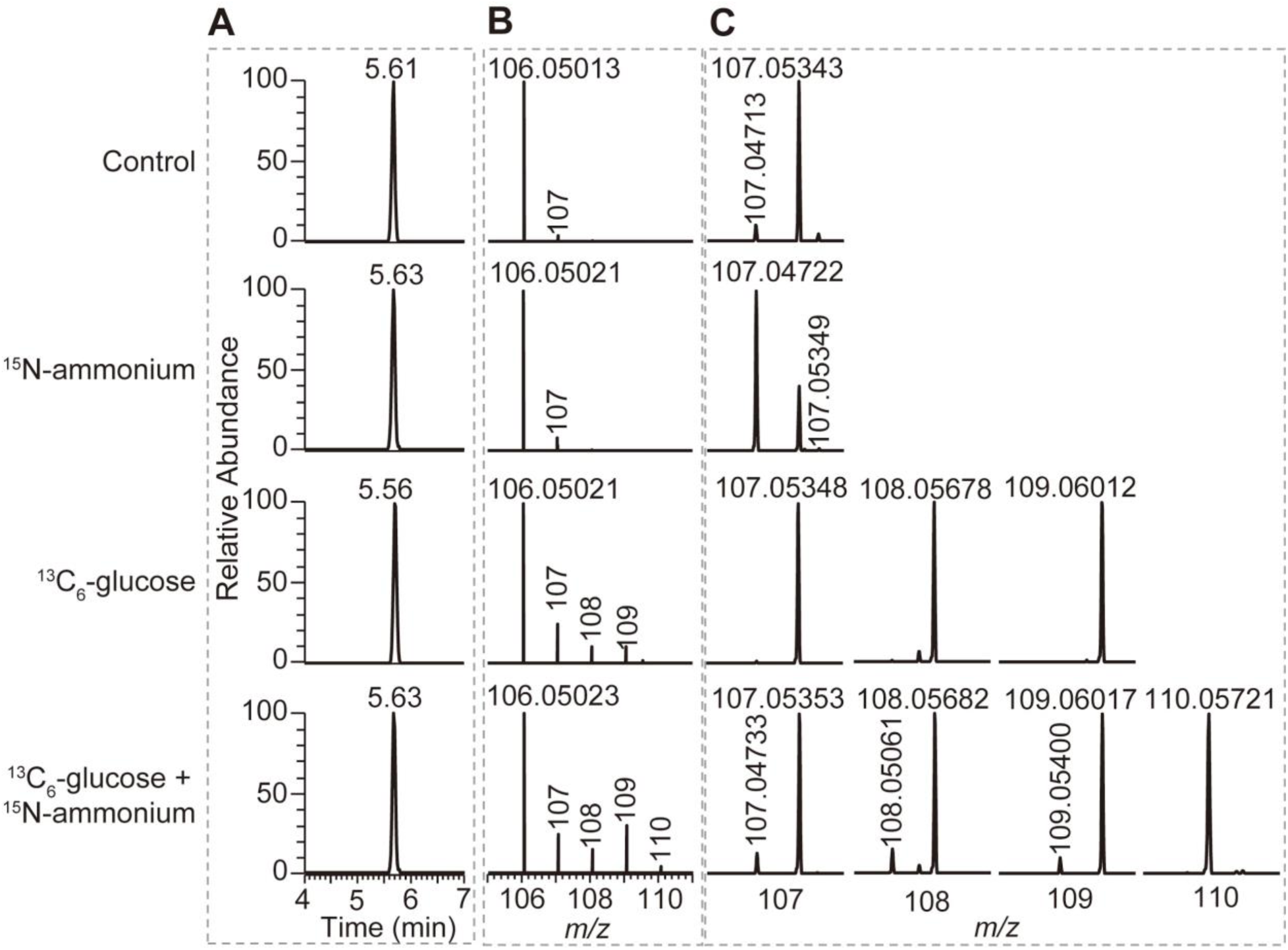
Identification of serine isolated from symbiotic *S. pistillata*. (**A, B**) Extracted ion chromatograms (EIC) (**A**) and the isotopic distributions (**B**) of serine isolated from symbiotic *S. pistillata* at different conditions. (**C**) The zoom of the area in (B) showing that different isotopologue compositions of serine are distinguished by HR-MS.

**Figure S19.**
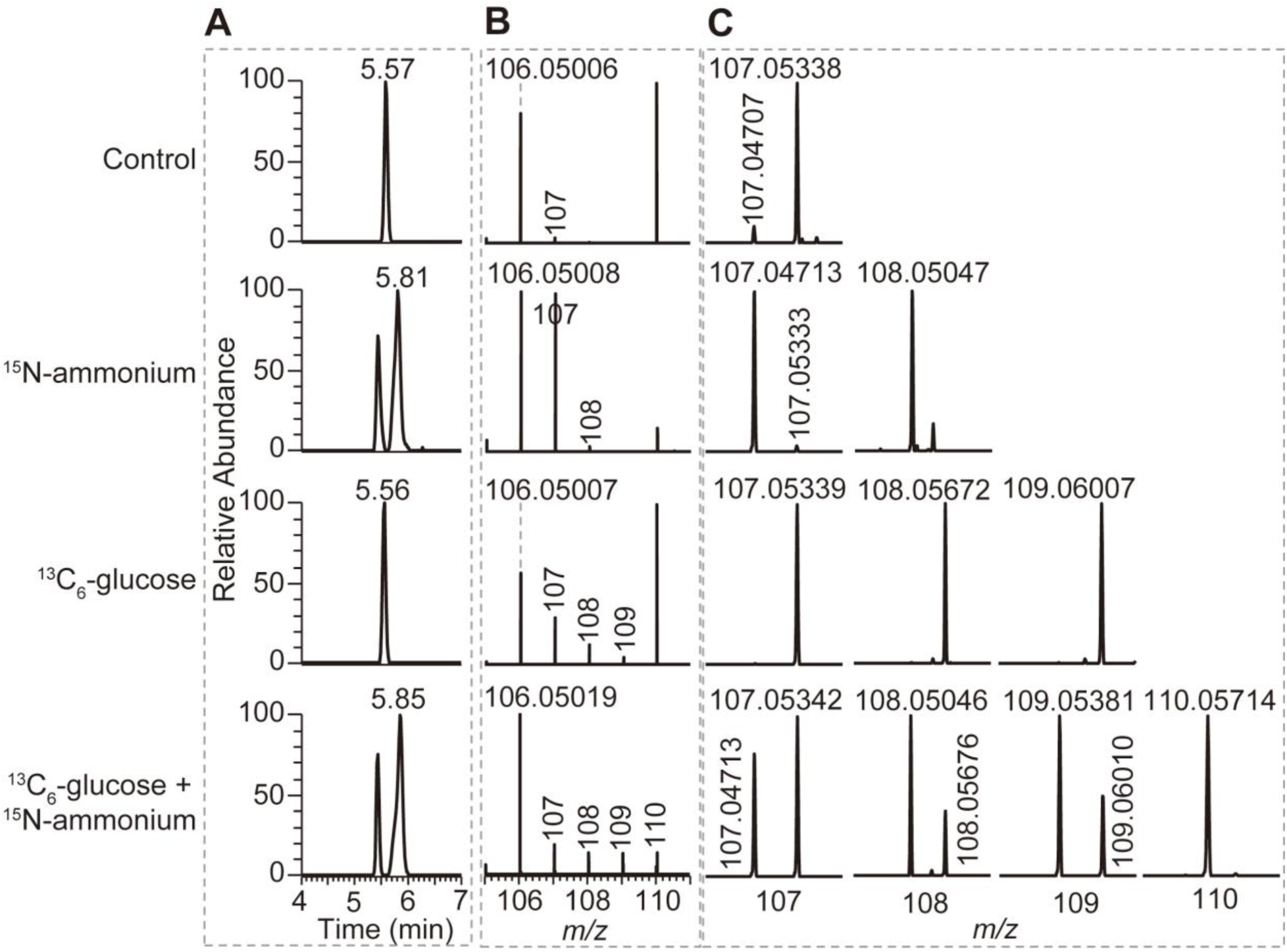
Identification of serine isolated from symbiotic *C. andromeda*. (**A, B**) Extracted ion chromatograms (EIC) (A) and the isotopic distributions (B) of serine isolated from symbiotic *C. andromeda* at different conditions. (**C**) The zoom of the area in (B) showing that different isotopologue compositions of serine are distinguished by HR-MS.

**Figure S20.**
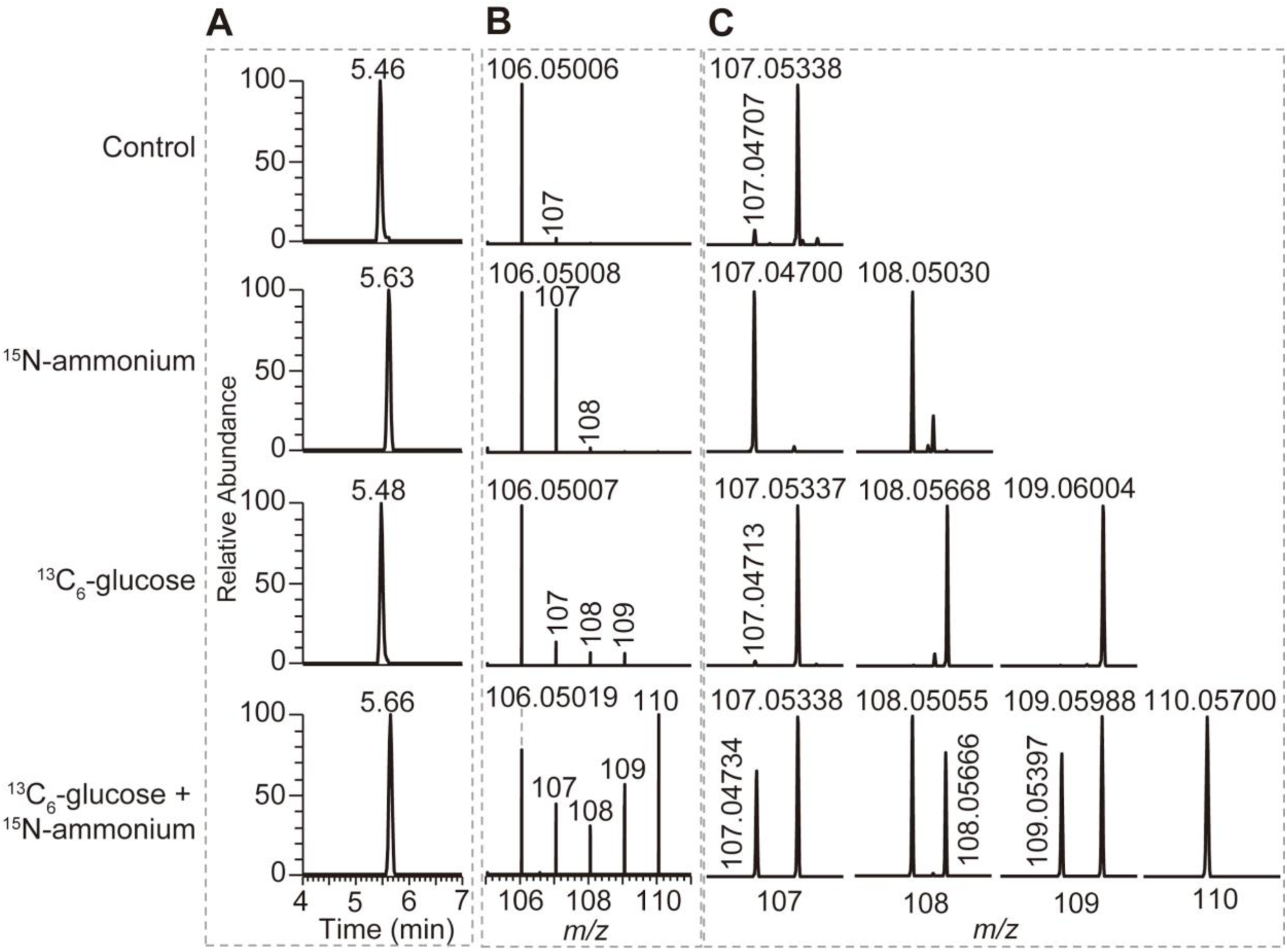
Identification of serine isolated from symbiotic *E. diaphana*. (**A, B**) Extracted ion chromatograms (EIC) (**A**) and the isotopic distributions (**B**) of serine isolated from symbiotic *E. diaphana* at different conditions. (**C**) The zoom of the area in (B) showing that different isotopologue compositions of serine are distinguished by HR-MS.

**Figure S21.**
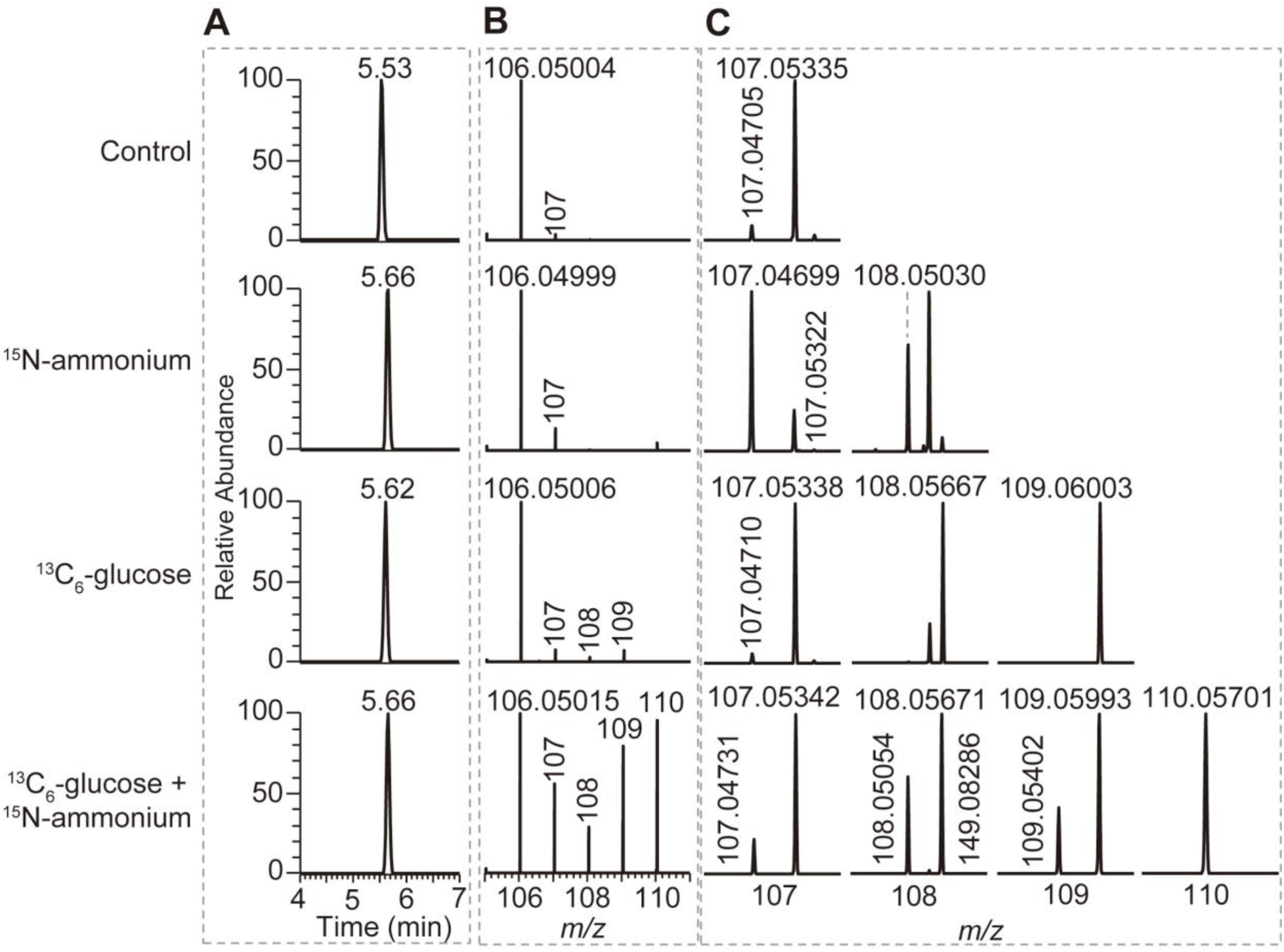
Identification of serine isolated from aposymbiotic *E. diaphana*. (**A, B**) Extracted ion chromatograms (EIC) (**A**) and the isotopic distributions (**B**) of serine isolated from aposymbiotic *E. diaphana* at different conditions. (**C**) The zoom of the area in (B) showing that different isotopologue compositions of serine are distinguished by HR-MS.

**Figure S22.**
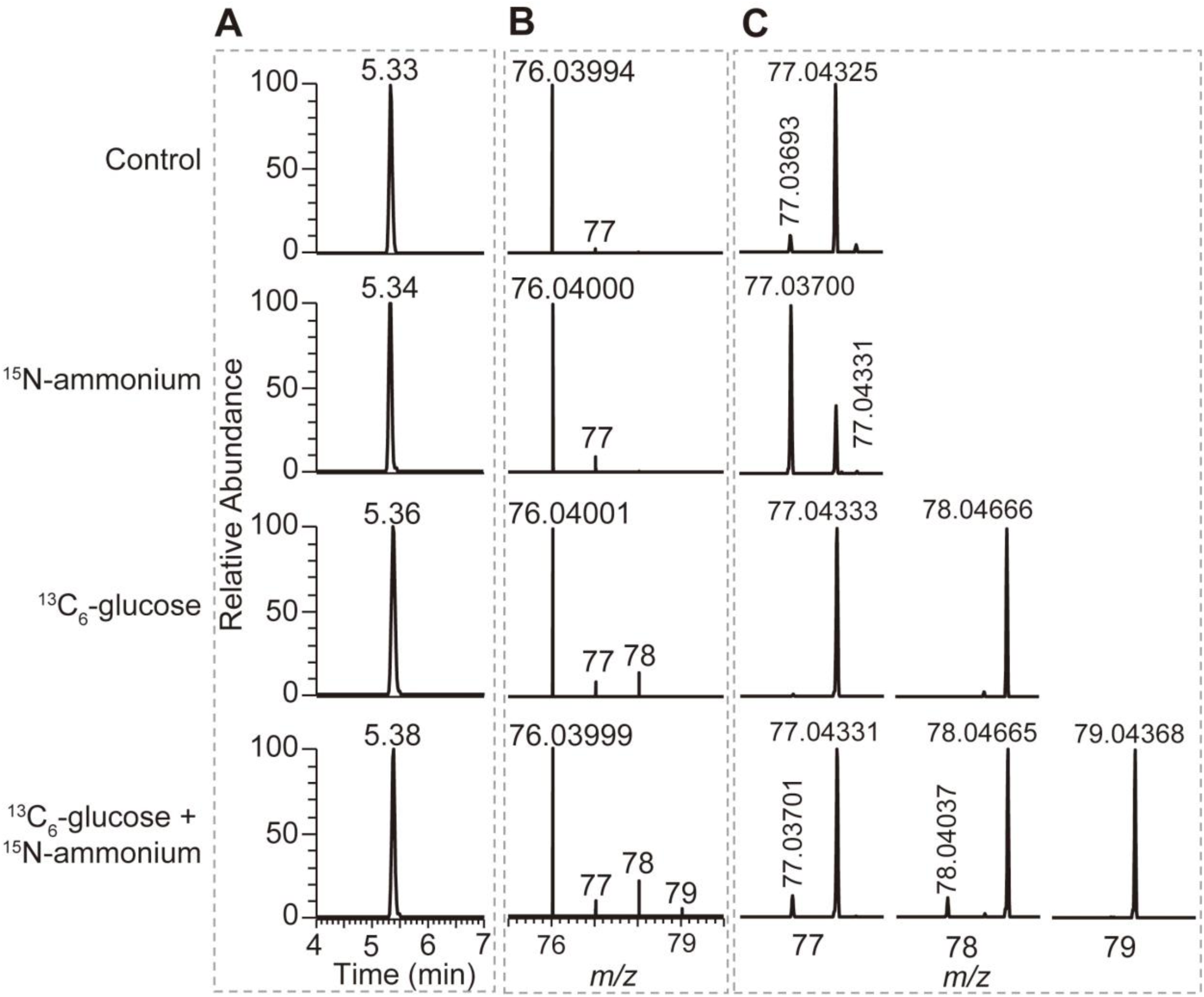
Identification of glycine isolated from symbiotic *S. pistillata*. (**A, B**) Extracted ion chromatograms (EIC) (**A**) and the isotopic distributions (**B**) of glycine isolated from symbiotic *S. pistillata* at different conditions. (**C**) The zoom of the area in (B) showing that different isotopologue compositions of glycine are distinguished by HR-MS.

**Figure S23.**
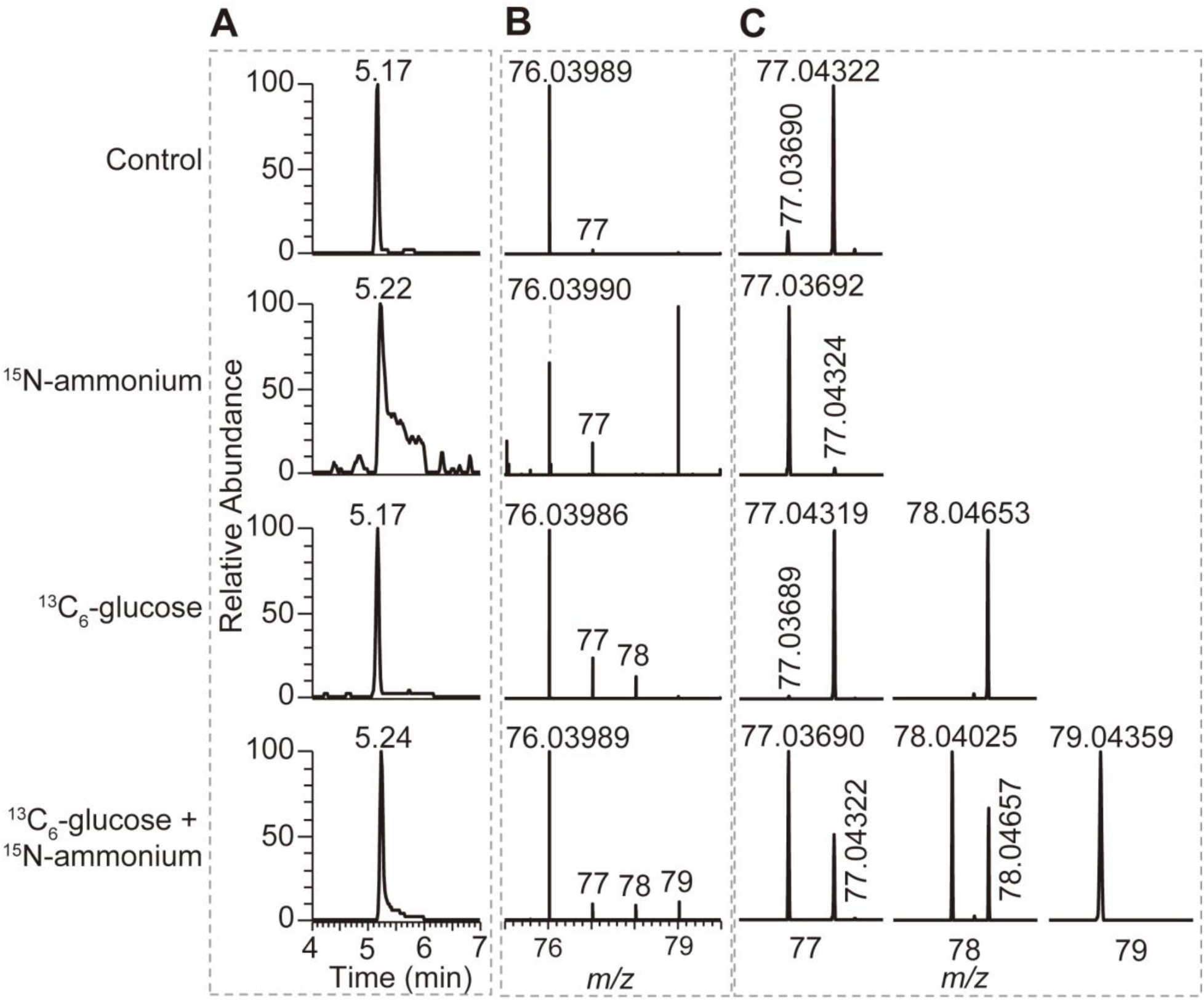
Identification of glycine isolated from symbiotic *C. andromeda*. (**A, B**) Extracted ion chromatograms (EIC) (**A**) and the isotopic distributions (**B**) of glycine isolated from symbiotic *C. andromeda* at different conditions. (**C**) The zoom of the area in (B) showing that different isotopologue compositions of glycine are distinguished by HR-MS.

**Figure S24.**
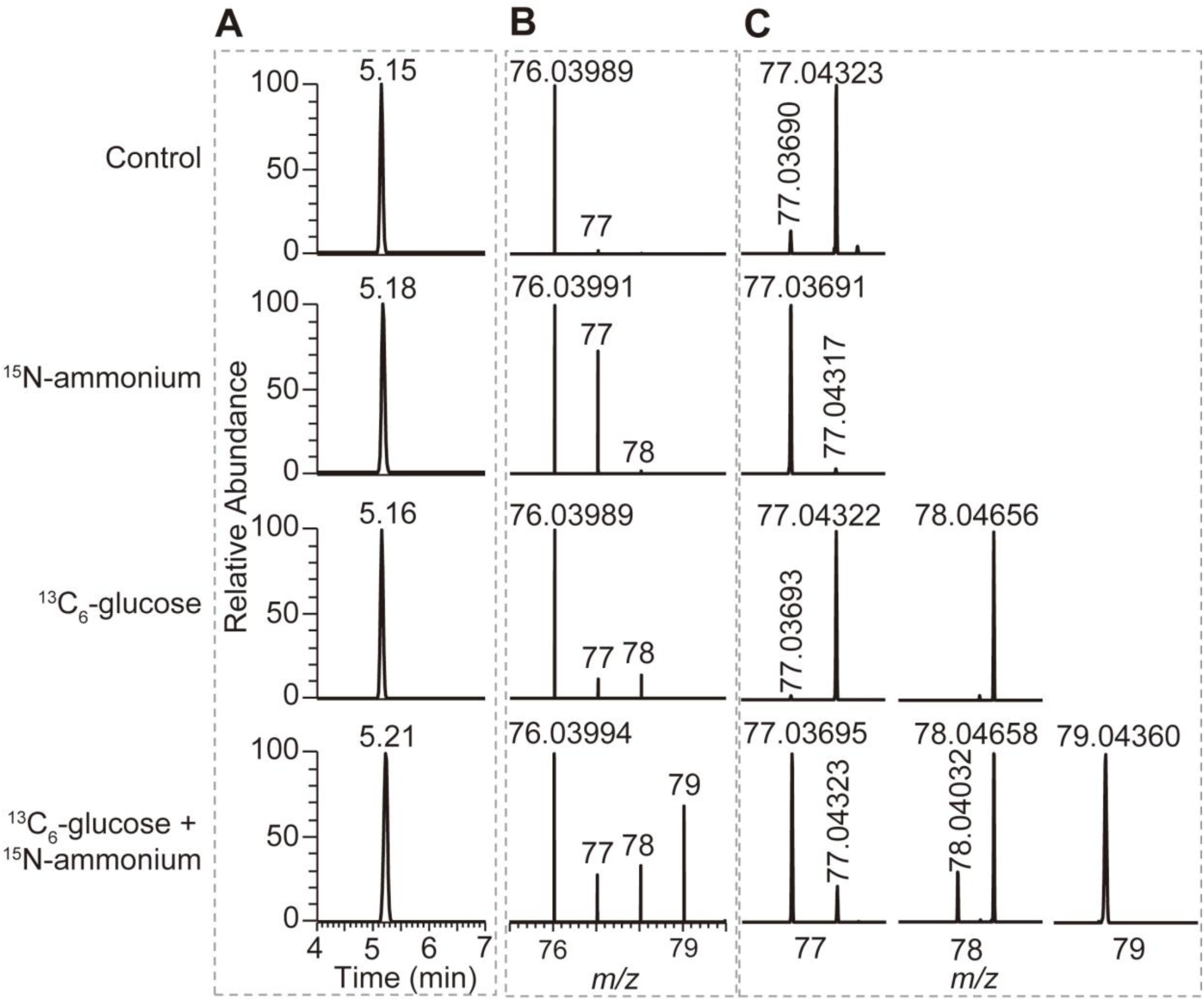
Identification of glycine isolated from symbiotic *E. diaphana*. (**A, B**) Extracted ion chromatograms (EIC) (**A**) and the isotopic distributions (**B**) of glycine isolated from symbiotic *E. diaphana* at different conditions. (**C**) The zoom of the area in (B) showing that different isotopologue compositions of glycine are distinguished by HR-MS.

**Figure S25.**
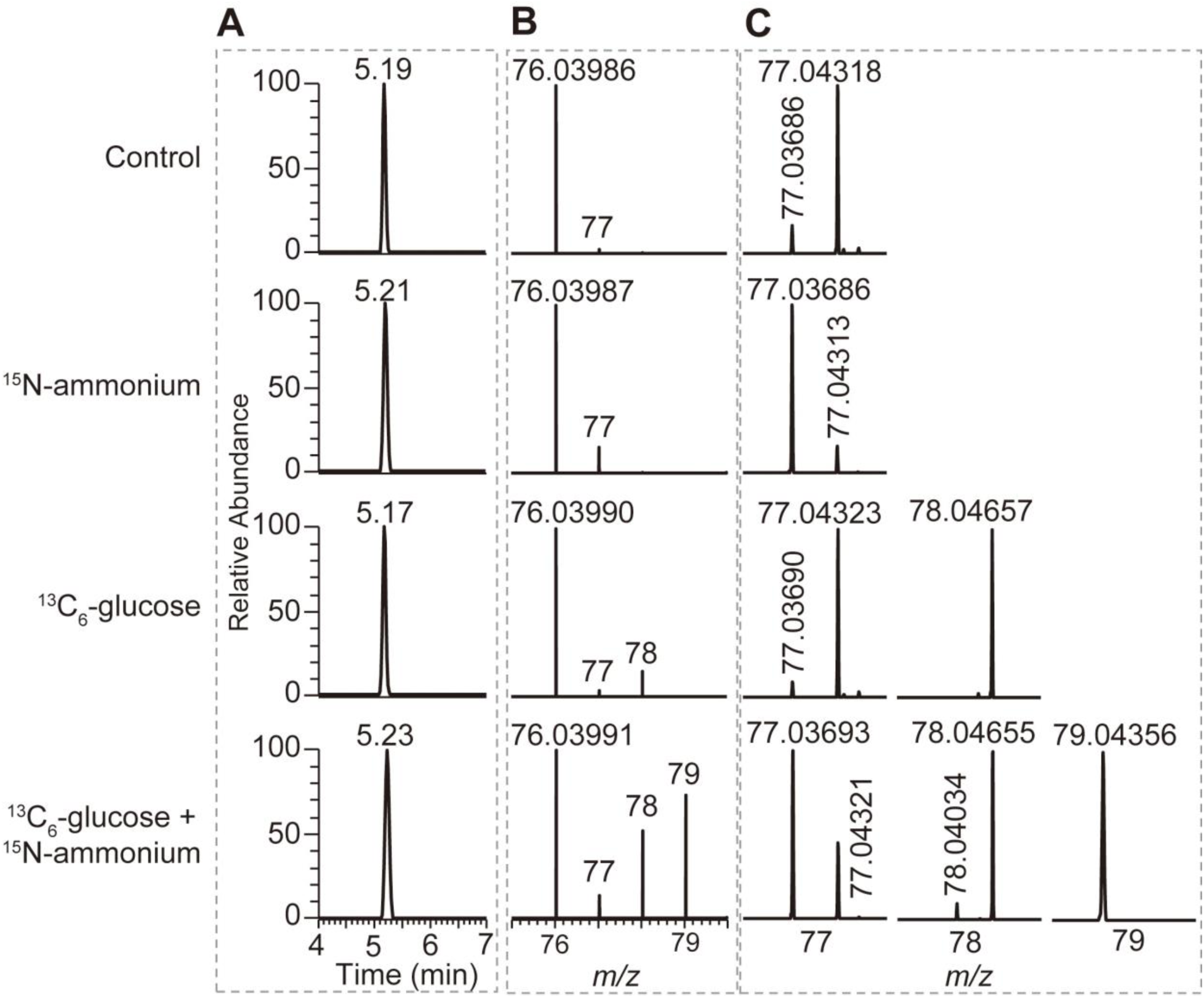
Identification of glycine isolated from aposymbiotic *E. diaphana*. (**A, B**) Extracted ion chromatograms (EIC) (**A**) and the isotopic distributions (**B**) of glycine isolated from aposymbiotic *E. diaphana* at different conditions. (**C**) The zoom of the area in (B) showing that different isotopologue compositions of glycine are distinguished by HR-MS.

**Figure S26.**
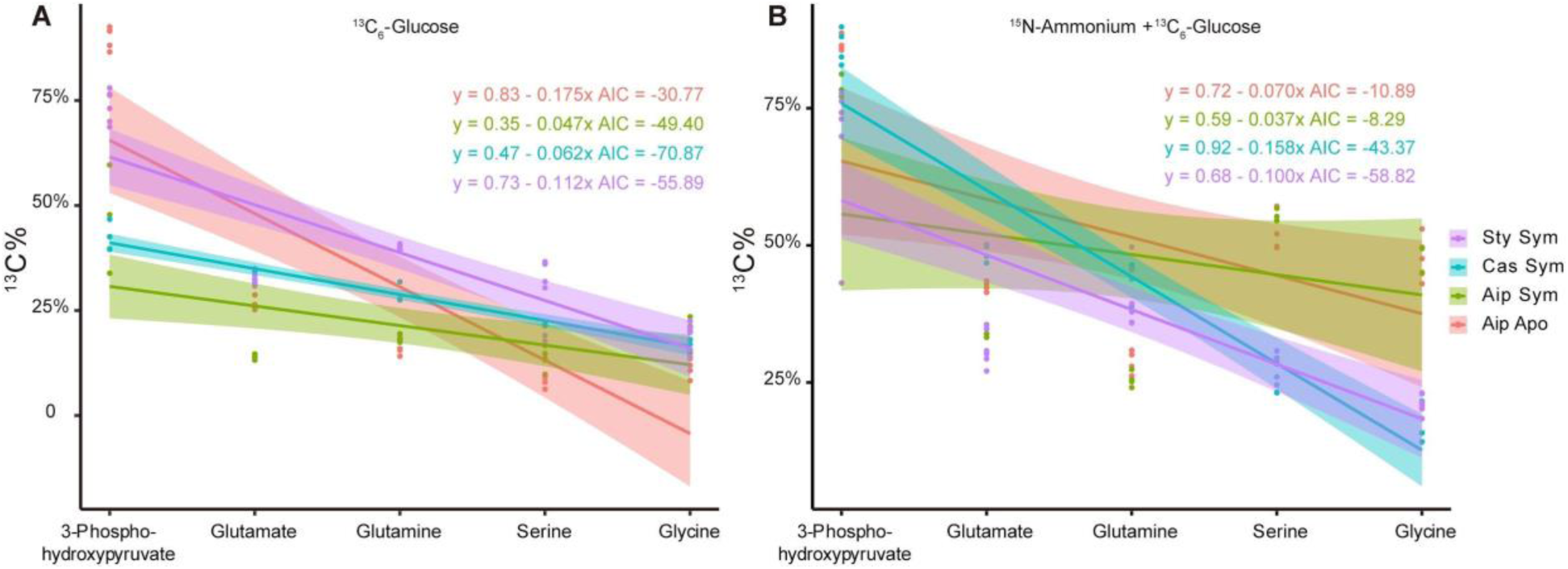
Generalized linear model of ^13^C proportion in metabolites across the GS/GOGAT-mediated amino acid synthesis pathway.

**Table S1.** Quantitative results of labeled metabolites isolated from symbiotic *S. pistillata*

**Table S2.** Quantitative results of labeled metabolites isolated from symbiotic *E. diaphana*

**Table S3.** Quantitative results of labeled metabolites isolated from symbiotic *C. andromeda*

**Table S4.** Quantitative results of labeled metabolites isolated from aposymbiotic *E. diaphana*

## Notes

### Competing Interest Statement

The authors have declared no competing interest.

